# Large-scale identification of ubiquitination sites on membrane-associated proteins in *Arabidopsis thaliana* seedlings

**DOI:** 10.1101/2020.09.16.299883

**Authors:** Lauren E. Grubb, Paul Derbyshire, Katherine Dunning, Cyril Zipfel, Frank L.H. Menke, Jacqueline Monaghan

## Abstract

Protein phosphorylation and ubiquitination are two of the most abundant forms of post-translational modifications in eukaryotes, regulated by thousands of protein kinases, phosphatases, E3 ubiquitin ligases, and ubiquitin proteases. Although previous studies have catalogued several ubiquitinated proteins in plants (Walton et al., 2016), few membrane-localized proteins have been identified. Receptor kinases (RKs) initiate phosphorylation signal relays that regulate plant growth, development, and stress responses. While the regulatory role of phosphorylation on protein kinase function is well-documented (Couto and Zipfel, 2016), considerably less is known about the role of ubiquitination on protein kinase function, even though protein turnover is critical to their signaling competence and cellular homeostasis. Here we describe the large-scale identification of ubiquitination sites on Arabidopsis proteins associated with or integral to the plasma membrane, including over 100 protein kinases.

## Introduction

Proteins can be mono-, poly-, and/or multi-mono-ubiquitinated, each affecting protein function in different ways. For example, mono-ubiquitination is often associated with protein activation or endocytosis, whereas poly-ubiquitination is often a mark for proteasomal degradation (Vierstra, 2012; Paez Valencia et al., 2016). Dynamic interplay between phosphorylation and ubiquitination has been observed in several proteins involved in immune signaling (Mithoe and Menke, 2018), including layered post-translational regulation of the receptor-like cytoplasmic kinase (RLCK) BIK1. BIK1 is directly phosphorylated and activated by several ligand-bound RKs (Couto and Zipfel, 2016), and can be dephosphorylated by the phosphatase PP2C38 (Couto et al., 2016). Precise control of BIK1 abundance is regulated by poly-ubiquitination by the E3 ligases PUB25 and PUB26 (Wang et al., 2018), as well as phosphorylation by the calcium-dependent protein kinase CPK28 (Monaghan et al., 2014; Wang et al., 2018) and the mitogen-activated protein kinase kinase kinase kinase (MAP4K) SIK1/MAP4K4 (Zhang et al., 2018; Jiang et al., 2019). Most recently, it was shown that BIK1 is also mono-ubiquitinated by the E3 ligases RHA3A and RHA3AB to regulate its activation and endocytosis (Ma et al., 2020).

## Results and Discussion

Proteomics and mutagenesis approaches have resulted in the discovery of several phosphorylated residues on BIK1 (Liang and Zhou, 2018). To help us understand the role of ubiquitination on BIK1 function, we set out to identify *in vivo* ubiquitination sites on BIK1. We enriched for plasma membrane-localized BIK1 by isolating microsomal protein fractions from Col-0/*pBIK1:BIK1-HA, cpk28-1/pBIK1:BIK1-HA* and *CPK28-OE1/pBIK1:BIK1-HA* genotypes, which express 100-fold higher levels of *BIK1* and differentially accumulate BIK1 protein compared to wild-type (Monaghan et al., 2014). To increase protein abundance and allow us to potentially capture immune-induced ubiquitination, proteasomal machinery was inhibited with 50 μM MG-132 an hour before treatment with water or 1 μM elf18 (an immunogenic peptide derived from bacterial EF-Tu (Zipfel et al., 2006)). Microsomal protein fractions were digested with trypsin, and anti-K-ε-GG agarose beads (Udeshi et al., 2013) were used to enrich ubiquitinated peptides by affinity binding. Ubiquitinated lysines were identified based on a shift of ∼114 Da -the mass of two glycine remnants that remain covalently bound to lysines following trypsin digestion -using liquid chromatography followed by tandem mass spectrometry (LC-MS/MS) (Supplementary Methods).

We confidently identified a total of 916 ubiquitinated peptides on 450 proteins across several biological replicates with a peptide false discovery rate of 0.025 (Table S1), and an additional 526 peptides on 398 proteins observed in single experiments (Table S2). Included in these data were seven ubiquitinated lysines on BIK1 (Tables 1, S1, S2, and Figure 1). Given our particular interest in BIK1, we manually inspected all spectra mapping to BIK1 and found an additional three sites (Figures 1 and S1), altogether corroborating five of the ubiquitinated residues reported by (Ma et al., 2020) and revealing five novel ones (Figure 1). Thus, BIK1 is ubiquitinated on multiple surface-exposed lysines *in vivo*: three in the N-terminal variable domain (K31, K41, K61), seven in the canonical kinase domain (K95, K106, K155, K170, K186, K286, K337), and five in the C-terminal region (K358, K366, K369, K374, K388) (Figure 1). Whether RHA3A/B and PUB25/26 compete for these sites or ubiquitinate distinct lysines remains to be tested experimentally, as does clarifying which E2 conjugating enzymes work with respective E3 ligases to catalyze these events (Turek et al., 2018). Furthermore, as the phospho-status of BIK1 has been shown to affect its regulation by both RHA3A/B and PUB25/26 (Wang et al., 2018; Ma et al., 2020), another challenge will be resolving the biochemical mechanisms underlying this interplay.

**Table 1:**
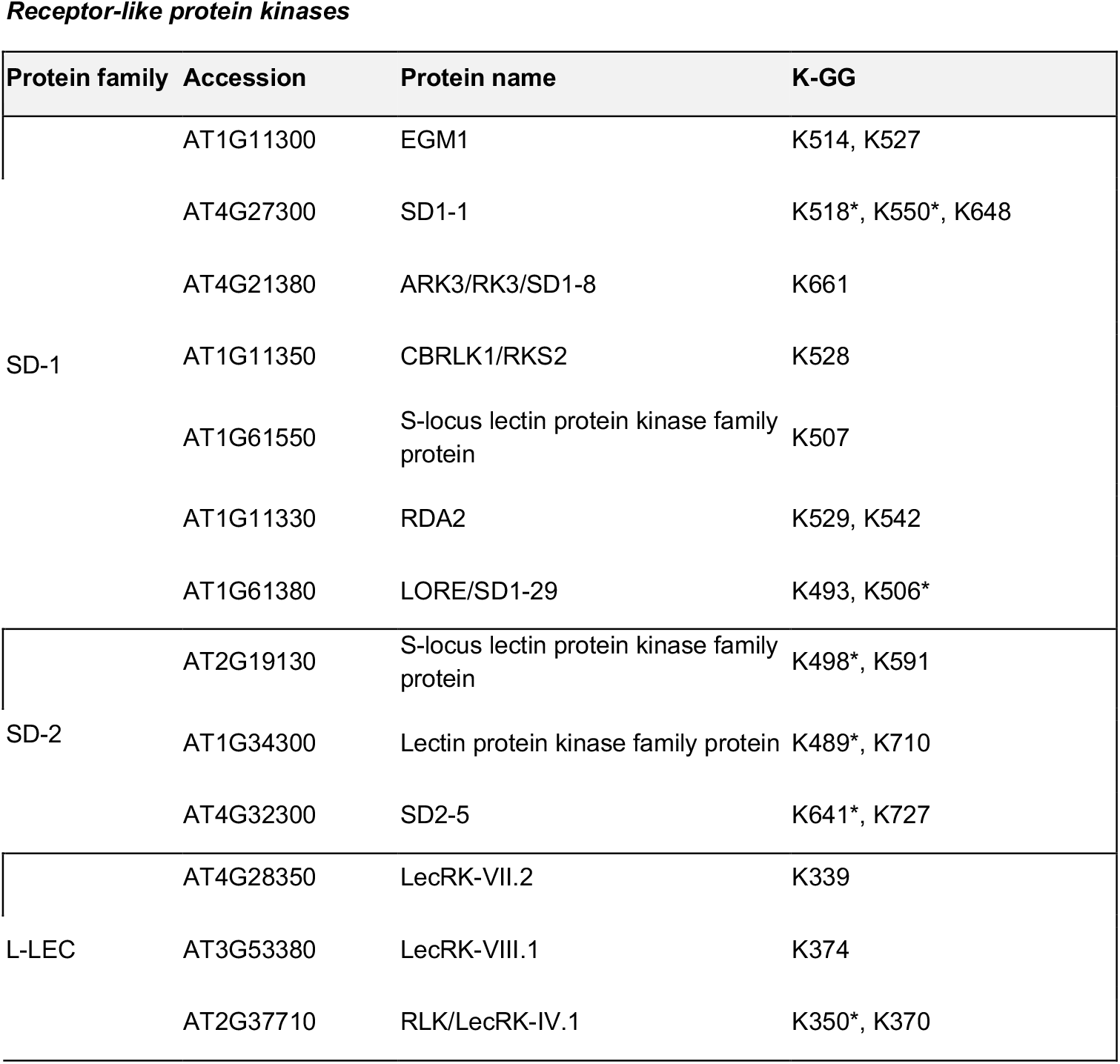
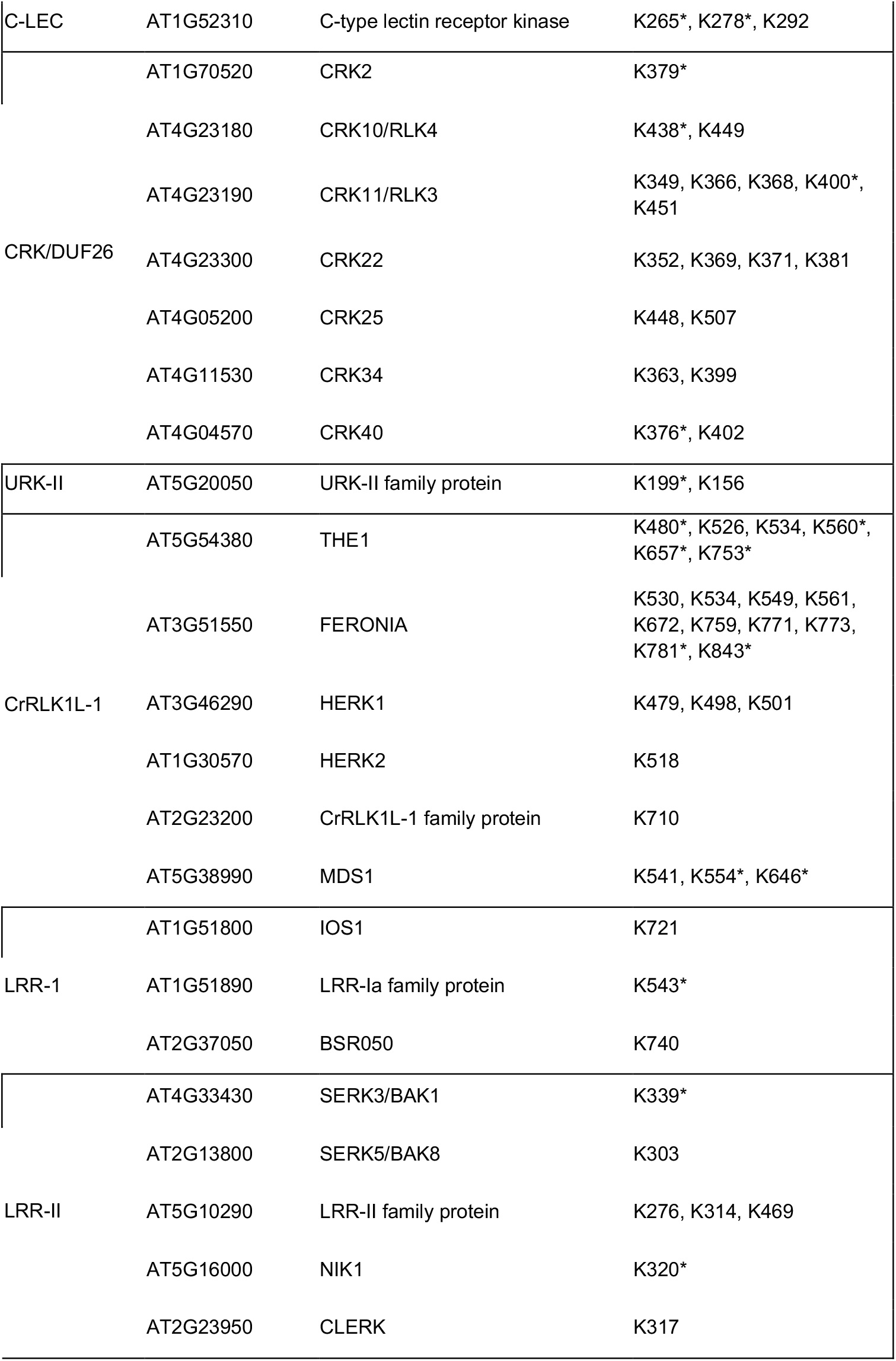
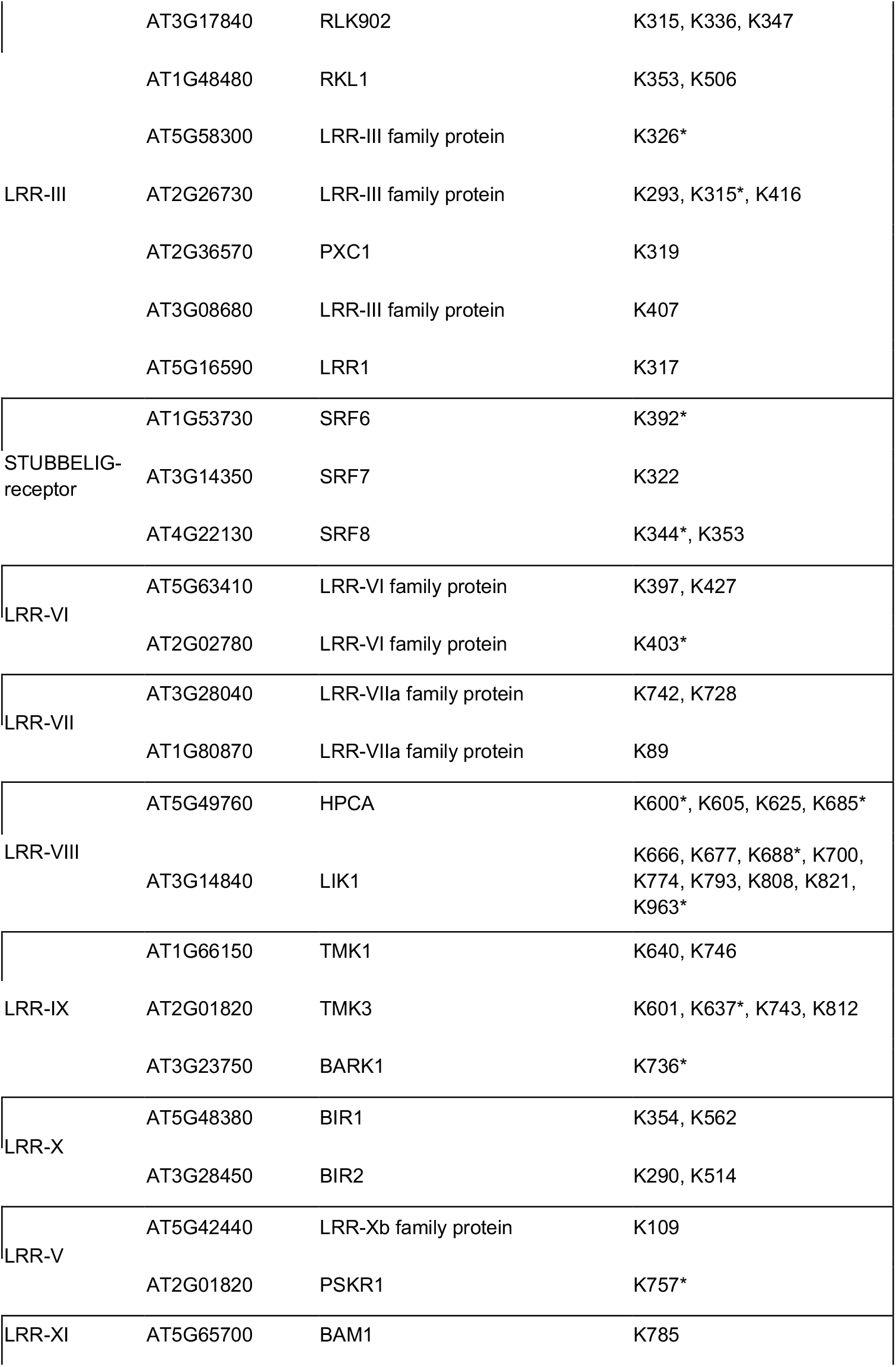
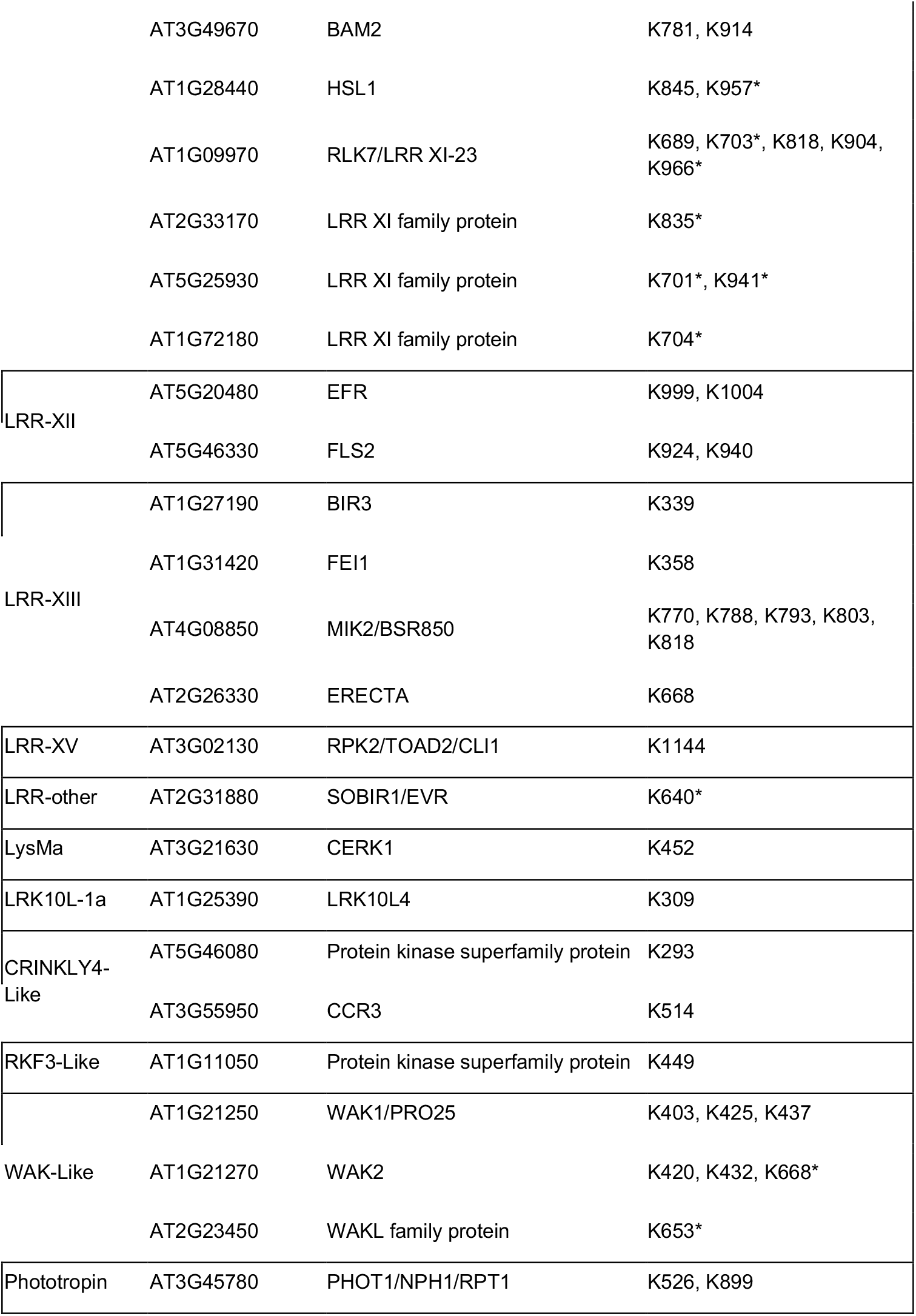
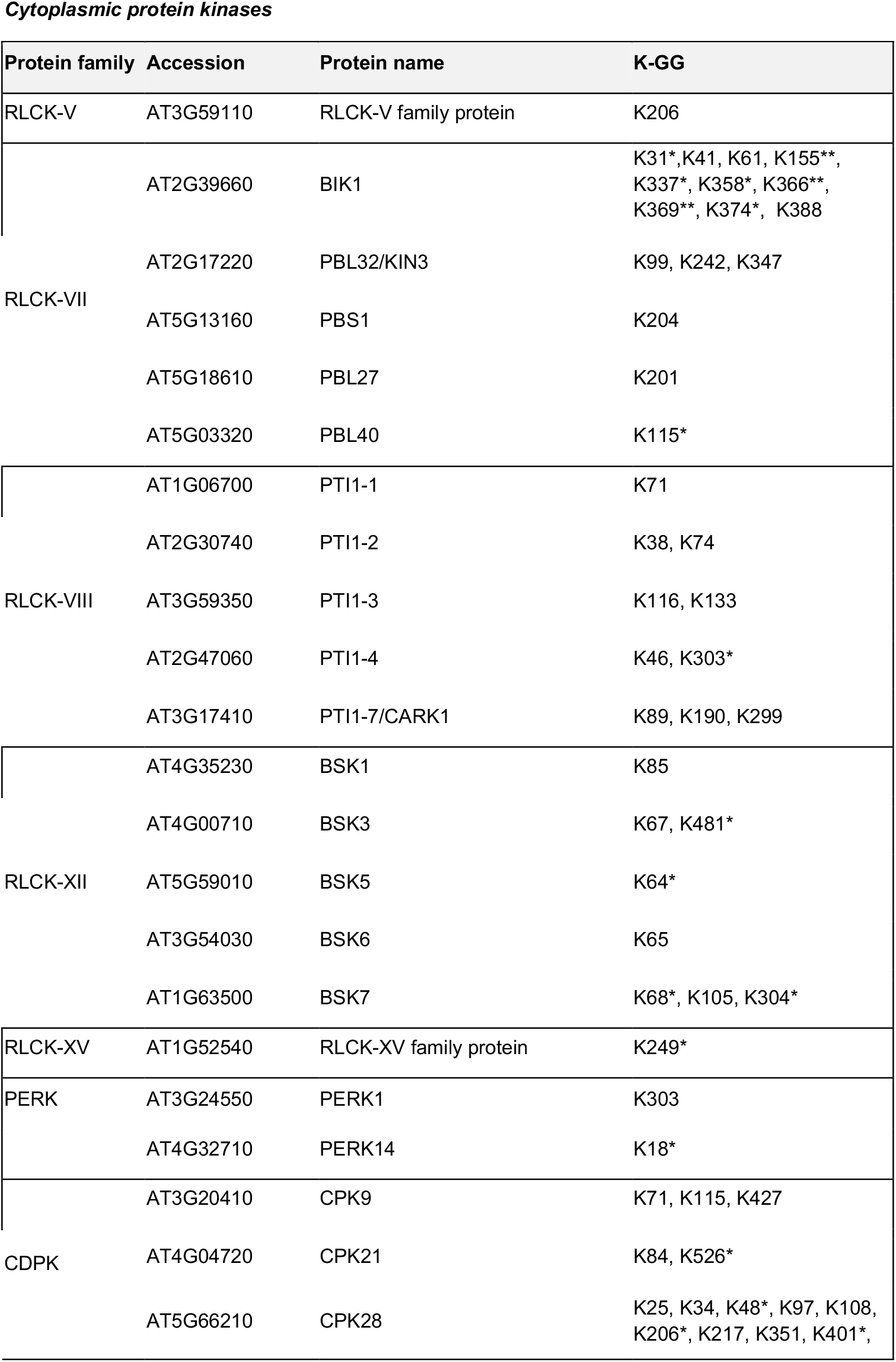
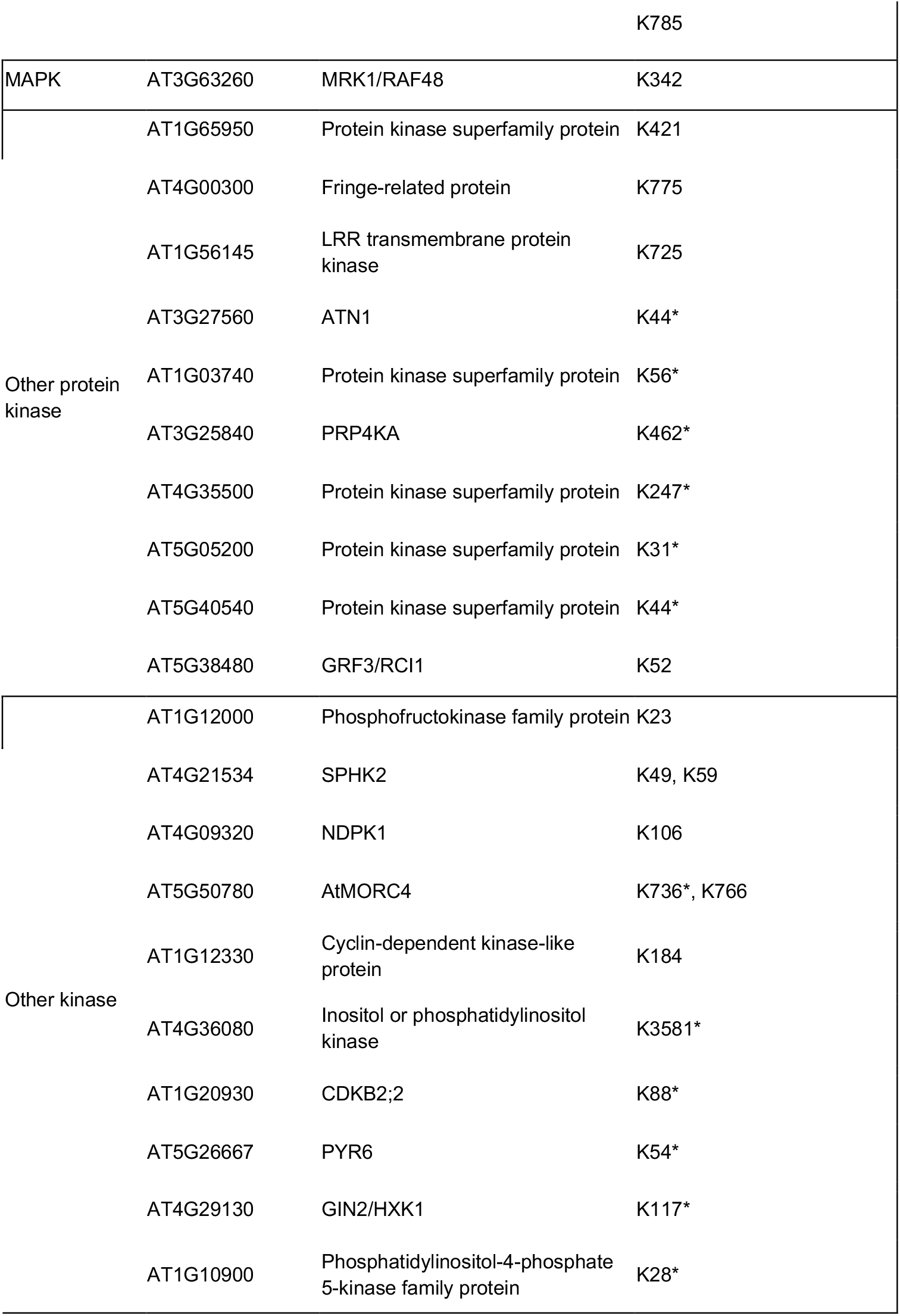
Ubiquitinated protein kinases identified in this study. Proteins matching the gene ontology term “kinase activity” were filtered from Supplementary Tables S1 and S2 and classified based on phylogenies presented by (Shiu and Bleecker, 2001; Shiu and Bleecker, 2003). *Residues that are only supported by a single observation (Table S2) are indicated by an asterisk and should be interpreted with caution. **Residues that were observed only after manual inspection of mass spectra matching BIK1 are indicated with two asterisks and shown in Supplementary Figure S1.

**Figure 1:BIK1 is ubiquitinated on multiple lysines *in vivo***.A comparison between this study and (Ma et al., 2020) indicates that BIK1 is ubiquitinated on three lysines at its amino (N) terminus, seven in its kinase domain, and five at its carboxyl (C) terminus. Ubiquitinated lysines identified in (Ma et al., 2020) are shown in green, those identified in this study are shown in blue, and residues identified in both studies are magenta. Although the structure of the BIK1 canonical kinase domain was recently solved (Lal et al., 2018), we modelled BIK1 in Phyre2 (Kelley et al., 2015) in order to include the disordered N- and C-terminal ends of the protein so that all of the identified sites could be shown in this surface representation of BIK1 in PyMol.

**Figure 1:**
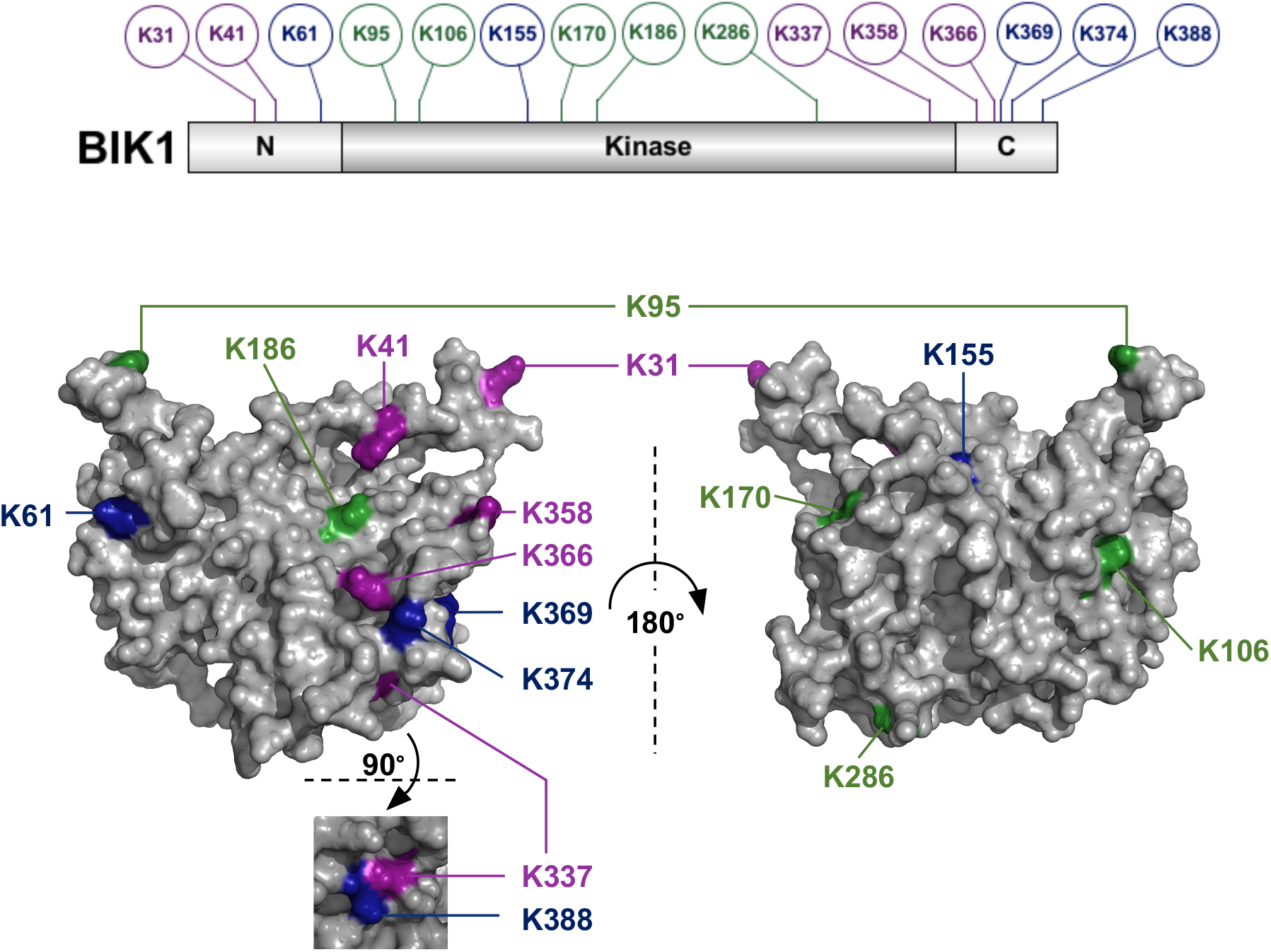
BIK1 is ubiquitinated on multiple lysines *in vivo*. A comparison between this study and (Ma et al., 2020) indicates that BIK1 is ubiquitinated on three lysines at its amino (N) terminus, seven in its kinase domain, and five at its carboxyl (C) terminus. Ubiquitinated lysines identified in (Ma et al., 2020) are shown in green; those identified in this study are shown in blue, and residues identified in both studies are magenta. Although the structure of the BIK1 canonical kinase domain was recently solved (Lal et al., 2018), we modelled BIK1 in Phyre2 (Kelley et al., 2015) in order to include the disordered N- and C-terminal ends of the protein so that we could include all of the identified sites in this surface representation of BIK1 in PyMol.

Analysis of gene ontology (GO) terms associated with proteins identified in the high-confidence dataset (Table S1) indicated an enrichment of proteins localized to the ‘plasma membrane’ (*p*=1.53×10^-114^) (Table S3). Because we analysed the samples in the mass spectrometer in data-dependent mode, without quantification, we are unable to comment on differences between genotypes or immune treatments. Therefore, any immune-triggered events must be corroborated experimentally. Multiple sequence alignments of peptides spanning -10 to +10 amino- and carboxyl-terminal to the modified lysines indicated very little consensus and no significant motifs (Figure S2). Unlike other post-translational modifications, the ubiquitination reaction requires coordination between E1 activating, E2 conjugating, and E3 ligase enzymes (Vierstra, 2012). While it may be possible for individual E2-E3 pairs to exhibit residue-level specificity on their target proteins, data from multiple species suggests that surface-availability may be the only unifying feature of ubiquitinated residues (Danielsen et al., 2011).

We identified ubiquitinated peptides mapping to proteins from diverse families, including aquaporins, H^+^ and Ca^2+^ ATPases, remorins, several classes of transporters, cellulose synthases, and others (Tables S1 and S2). Comparison between our dataset and 8 published Arabidopsis ubiquitome datasets, as well as manual inspection of the literature, revealed 265 novel ubiquitin targets (Table S4). We noted that molecular function GO terms ‘protein modification’ (*p*=1.79×10^-12^), ‘phosphorylation’ (*p*=2.15×10^-26^), and ‘response to stimulus’ (*p*=6.44×10^-21^) were particularly enriched in our dataset (Table S3). Interestingly, we identified multiple ubiquitinated lysines on over 70 RKs representing diverse subgroups, including FLS2, EFR, CERK1, LORE, RLK7, SOBIR1/EVR, LIK1, RKL1, WAK1, WAK2, FER, ER, BAM1, BAM2, and others (Table 1). We also identified ubiquitination sites on more than 20 plasma membrane-associated cytoplasmic protein kinases from several subgroups (Table 1). Because analysis of tryptic peptides with ubiquitinated lysine residues enriched by anti-K-ε-GG does not allow for discrimination between mono- or poly-ubiquitination, it is likely that we have captured both degradative and non-degradative ubiquitination on these protein kinases. Given the broad interest in phosphorylation-based signal transduction and protein homeostasis, we expect this information will be valuable to the plant research community, and look forward to future studies that explore the function of these ubiquitination events.

## Footnotes

## Acknowledgements

We thank Jan Sklenar for helpful suggestions and technical assistance, and are grateful to Melissa Bredow for help using Phyre2 and PyMol. We thank Libo Shan and Ping He for sharing data prior to publication.

## Author Contributions

F.L.H.M. and J.M. designed the research; L.E.G. and F.L.H.M. performed the experiments; L.E.G., P.D., K.D., F.L.H.M., and J.M. analyzed the data; C.Z., F.L.H.M., and J.M. supervised the work; J.M. wrote the letter with input from all authors.

## Supplemental Data

Grubb LE, Derbyshire P, Dunning K, Zipfel C, Menke FLH, and Monaghan J. “Large-scale identification of ubiquitination sites on membrane-associated proteins in *Arabidopsis thaliana* seedlings”

## Supplemental Methods

The plant genotypes used in this study have been previously described (Monaghan et al., 2014). Twenty mg of seed from each genotype was surface-sterilized using chlorine gas and stratified at 4°C in the dark for 2 days. Seed was then transferred to 250 mL Erlenmeyer flasks containing 50 mL 0.5X Murashige and Skoog (MS) media supplemented with 0.05% sucrose, and shaken at 100 rpm at ambient temperature with a 10h photoperiod for 8 days. Seedlings were incubated with 50 μM MG-132 (Sigma Aldrich, UK) for 1 h with shaking, and then vacuum infiltrated with water (mock) or 1 μM elf18 peptide (EZ Biolabs, USA) for 10 min. Samples were flash frozen and ground to a coarse powder with a mortar and pestle in liquid N_2_, then homogenized in urea lysis buffer (8 M urea, 50 mM Tris-HCl pH 8.0, 150 mM NaCl, 1 mM EDTA, 1 mM P9599 protease inhibitor (Sigma Alrich), 1 mM phenylmethylsulfonyl fluoride (PMSF), 50 μM PR-619 (Sigma Aldrich)) using equal volume-to-tissue ratio in a Potter tube for 10 min at 1,000 rpm. An aliquot of 1 mL was removed and kept for quick analysis of elf18-induced MAPK activation by immunoblot as previously described (Monaghan et al., 2014) (data not shown). The remaining homogenate was centrifuged at 5,856 x *g* for 1 h at 4°C. The supernatant was transferred to a polycarbonate tube and centrifuged at 110,000 x *g* for 1 h at 4°C. The pellet was resuspended in 2 mL of lysis buffer with PMSF, and protein concentration was determined using the Bradford Assay according to the manufacturer’s instructions (BioRad). Three mg of protein in 2 mL lysis buffer was incubated with 5 mM Tris(2-carboxyethyl)phosphine (TCEP) at ambient temperature for 45 min. Iodoacetamide (10mM) was added, and samples were incubated in the dark for 30 min at ambient temperature. Finally, samples were diluted to 10 mL in 50 mM Tris-HCl pH 8.0 and digested with 30 μg of trypsin (Pierce, UK) at 30°C overnight.

Following trypsin digestion, samples were acidified with aliquots of 50% trifluoroacetic acid (TFA) to pH 3.0. Precipitate was removed by centrifugation at 1,600 x *g*, and the peptides in the supernatant were purified using 2.5 mL C18 silica reversed-phase chromatography Sep-Pak columns (Waters). Prior to use, the columns were equilibrated sequentially with 1 column volume (CV) MeOH, then 1 CV activation buffer (80% acetonitrile (ACN), 0.1% TFA), and then 5 CV of wash buffer (2% ACN, 0.1% TFA). Sample was added to the column and allowed to drip by gravity flow, collected, and applied to the column a second time before application of 5 CV wash buffer. Trypsin-digested peptides were eluted in 2 CV elution buffer (40% ACN, 0.1% TFA) and lyophilized.

The PTMScan® Ubiquitin Remnant Motif K-ε-GG antibody (Cell Signaling Technologies, UK) was first crosslinked to the agarose beads by washing each tube 3 times with 1 mL of 100 mM sodium borate pH 9.0, with centrifugation at 2,000 x *g* between each wash. The beads were then resuspended in 20 mM dimethyl pimelimidate (DMP) crosslinker and 100 mM sodium borate, and incubated for 1 h at room temperature with end-over-end rotation. The beads were washed twice with 1 mL of 200 mM ethanolamine pH 8.0, resuspended in 200 mM ethanolamine, and incubated overnight at 4°C with end-over-end rotation.

Prior to immunoaffinity purification (IP), crosslinked beads were washed 3 times with 1.5 mL IAP buffer (50 mM MOPS pH 7.2, 10 mM sodium phosphate, 50 mM NaCl). Lyophilized peptide samples were resuspended in 1.5 mL IAP buffer, sonicated for 10 min, and centrifuged at 16,000 x *g* for 5 min. A volume of 1,300 μL of peptide sample was added to 300 μL of K-ε-GG beads in IAP buffer, and the remaining 200 μL was reserved as ‘input’ sample. The IP was conducted for 2 h at 4°C with end-over-end rotation, followed by centrifugation for 1 min at 2,000 x *g*. The supernatant was kept as the ‘unbound’ sample to test the efficiency of di-Gly enrichment. Beads were washed twice with 1.5 mL IAP buffer, then 3 times with high performance liquid chromatography (HPLC) grade water prior to elution with 100 μL of 0.15% TFA, and centrifugation 1 min at 2,000 x *g*. The supernatant was kept as the ‘elution’ sample for analysis by LC-MS/MS.

Input, unbound, and elution samples were partially purified using C18 Micro-Spin Stagetips columns (The Nest Group Inc.). The samples were acidified with 50% TFA to a pH of 3.0, and the columns were placed in 2 mL microcentrifuge tubes and equilibrated by washing 3 times with 200 μL MeOH and centrifuged at 135 x *g* for 30 s between each wash. Equilibration continued with 3 washes of 200 μL equilibration buffer (80% ACN, 0.1% TFA) and centrifugation for 2 min at 185 x *g*, followed by washing 6 times with wash buffer (2% ACN, 0.1% TFA). Peptide solutions were loaded twice into the equilibrated columns and centrifuged for 4 min at 240 x *g*. Columns were then washed 6 times with Wash Buffer as above, followed by tandem elutions in 150 μL elution buffer (40% ACN, 0.1% TFA).

To prepare for mass spectrometry, the input, unbound, and eluted samples were dehydrated using vacuum centrifugation (SpeedVac), and then resuspended in 2% ACN and 0.1% Formic Acid (FA), followed by vortexing and sonication for 10 min prior to a final centrifugation at 10,000 xg for 10 min. Samples were then loaded into a 96-well plate for analysis on an Orbitrap Fusion Mass Spectrometer (ThermoFisher).

LC-MS/MS analysis was performed as described in (Bender et al., 2017) with the following modifications. On the Orbitrap Fusion MS/MS spectra were triggered with data dependent acquisition method using ‘top 20’ and ‘most intense ion’ settings. Peak lists in Mascot generic file format (.mgf files) were generated from raw files by using the MSConvert package (Matrix Science). Peak lists were searched using Mascot server v.2.4.1 (Matrix Science) against TAIR database (version 10), a separate in-house constructs database, and an in-house contaminants database. Tryptic peptides with up to 2 possible mis-cleavages and charge states +2, +3, +4, were allowed in the search. The following modifications were included in the search: oxidized methionine, diGly on lysine as variable modification and carbamido-methylated cysteine as static modification. Data were searched with a monoisotopic precursor and fragment ions mass tolerance 10ppm and 0.6 Da respectively. Mascot results were combined in Scaffold v. 4 (Proteome Software) and exported in Excel (Microsoft Office).

## Supplemental Tables

**Table S1: High-confidence peptides identified in multiple experiments**.

Peptides shown were filtered for the ubiquitin remnant diGly and were observed at least twice with a minimal Mascot score of 20. The peptide level false discovery rate was 0.025. Columns J to AR report the number of spectra observed in total (J), subtotals per genotype and treatment (K-Q) or in individual samples (S-AR). Column I shows whether the assignment of the diGly modified residue is possibly ambiguous (P) or not ambiguous (N).

**Table S2: Peptides identified in single experiments**.

Peptides shown were filtered for the ubiquitin remnant diGly and were observed at once with a minimal Mascot score of 20. The peptide level false discovery rate was 0.025. Columns J to AR report the number of spectra observed in total (J), subtotals per genotype and treatment (K-Q) or in individual samples (S-AR). Column I shows whether the assignment of the diGly modified residue is possibly ambiguous (P) or not ambiguous (N).

**Table S3: Gene ontology terms associated with proteins identified in this study**.

Gene ontology profiling was conducted using unique protein identifiers from Table S1 in g:profiler (Raudvere et al., 2019).

**Table S4: Comparative analysis reveals 268 unique ubiquitin targets identified in this study**.

Ubiquitinated proteins identified from earlier large-scale studies (Maor et al., 2007; Manzano et al., 2008; Igawa et al., 2009; Kim et al., 2013; Svozil et al., 2014; Johnson and Vert, 2016; Walton et al., 2016; Romero-Barrios et al., 2020) were compared to those identified in this study (Table S1) using Microsoft Excel. Manual inspection of the literature revealed additional ubiquitinated proteins, as well as proteins predicted to be ubiquitinated.

## Supplemental Figures

**Figure S1:**
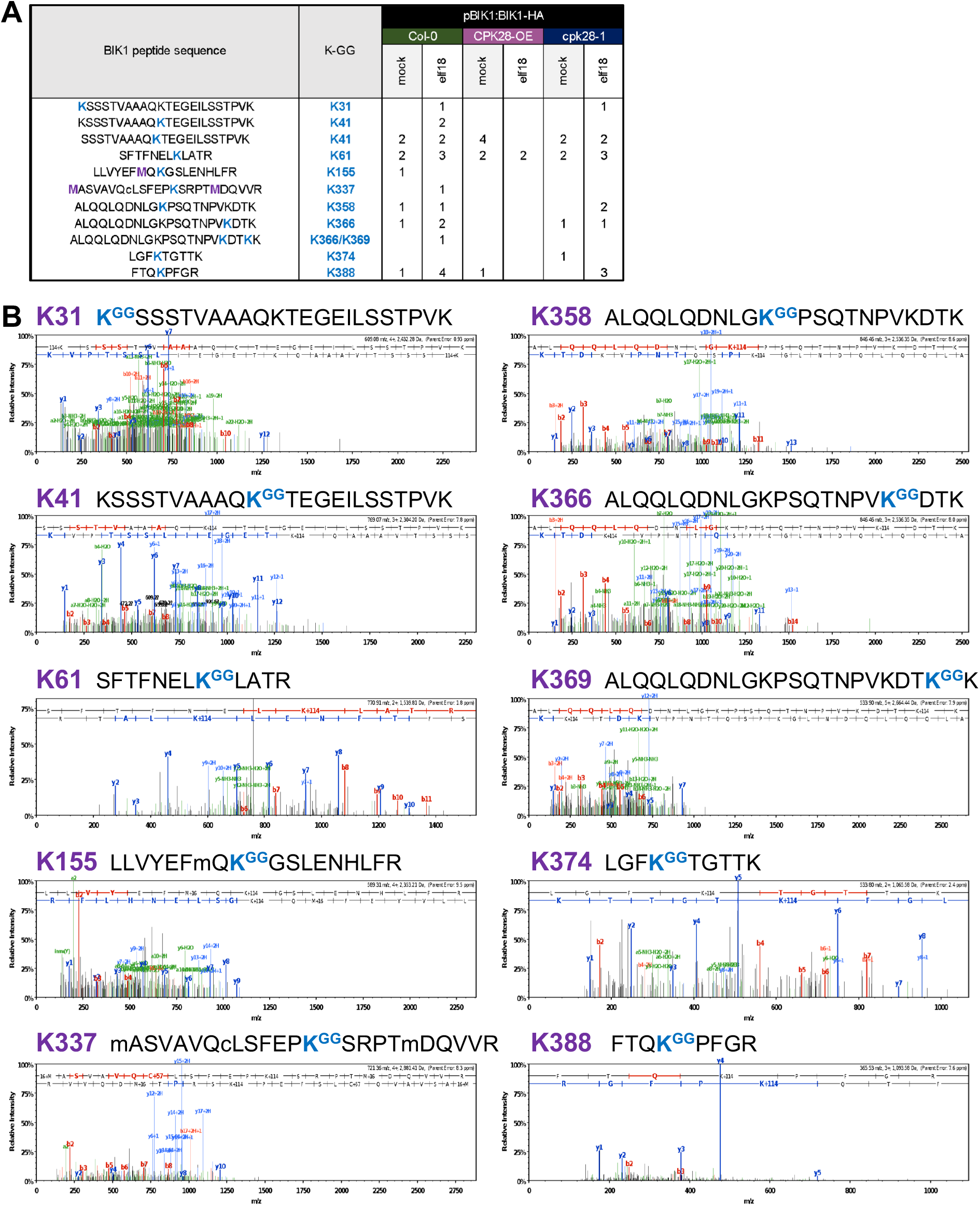
Ubiquitinated residues identified on BIK1. Ten tryptic peptides mapping to BIK1 contain di-Gly remnants in different genotypes and treatments as outlined in (A); the modified Lys residue is coloured blue. Mass spectra for each peptide were extracted from Scaffold and shown in (B); B- and Y-ions are coloured in red and blue, respectively.

**Figure S2:**
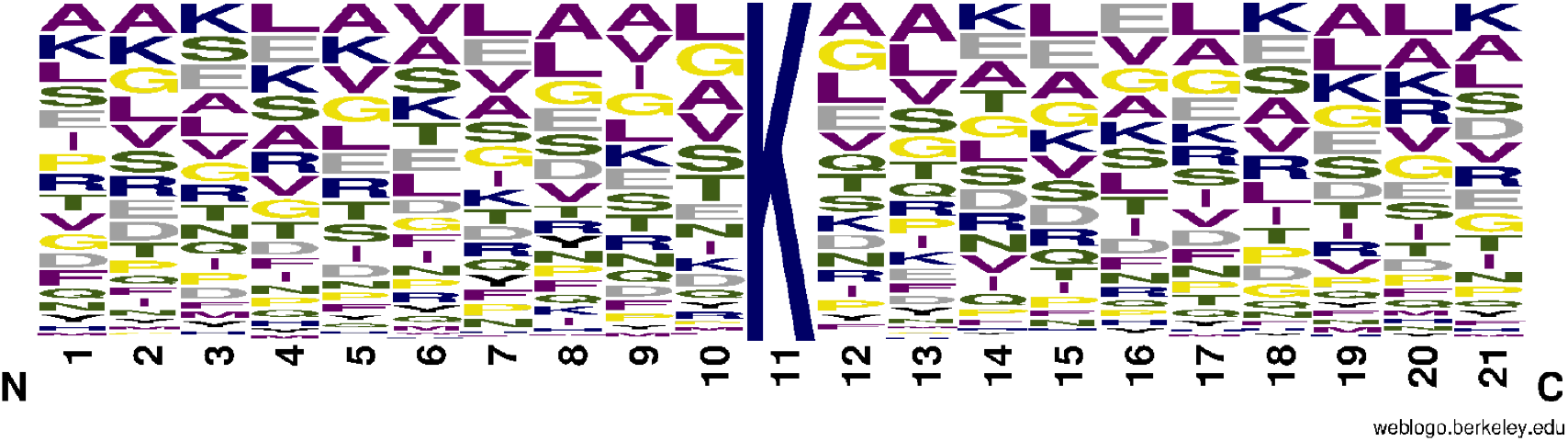
Consensus motif analysis of ubiquitinated lysines. Non-redundant full-length protein sequences were retrieved from The Arabidopsis Information Resource (TAIR) and peptide sequences -10 and +10 amino- and carboxyl-terminal to the modified lysines were identified using the MID function in Microsoft Excel. These 21-amino-acid peptide sequences were analyzed using the MEME Suite v5.1.1 MoMo motif finder tool (Bailey et al., 2009), however no statistically significant (*p*<0.05) consensus motifs were identified. A multiple-sequence alignment of the peptide sequences surrounding the modified lysine was created using WebLogo (Crooks et al., 2004). Blue residues contain positively-charged R-groups (KHR); Black residues are negatively-charged (DE); Magenta residues are hydrophobic (AILMFWYV); Green residues are polar uncharged (STNQ); Yellow residues are special cases (CGP).

**Table S4.**
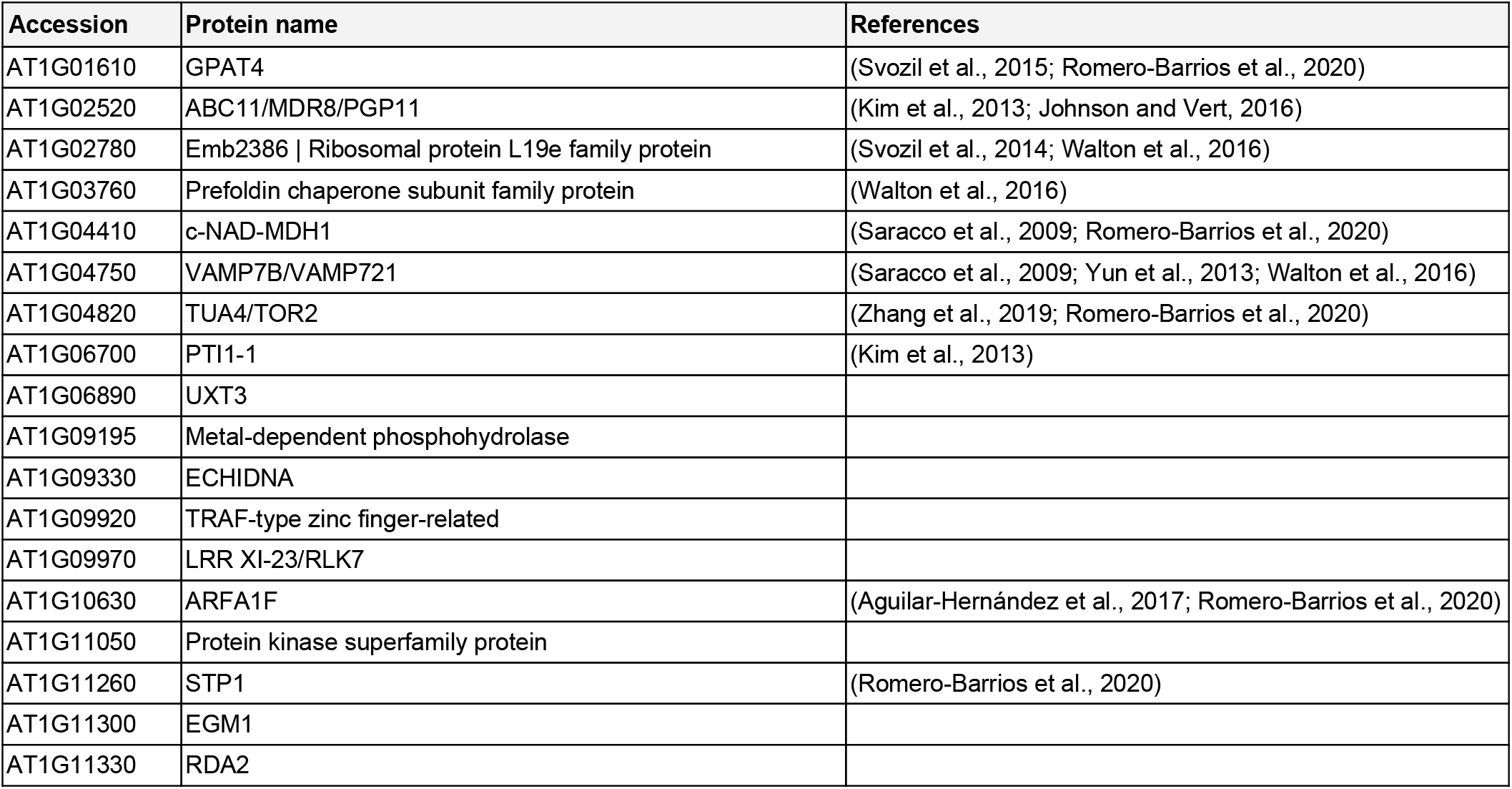
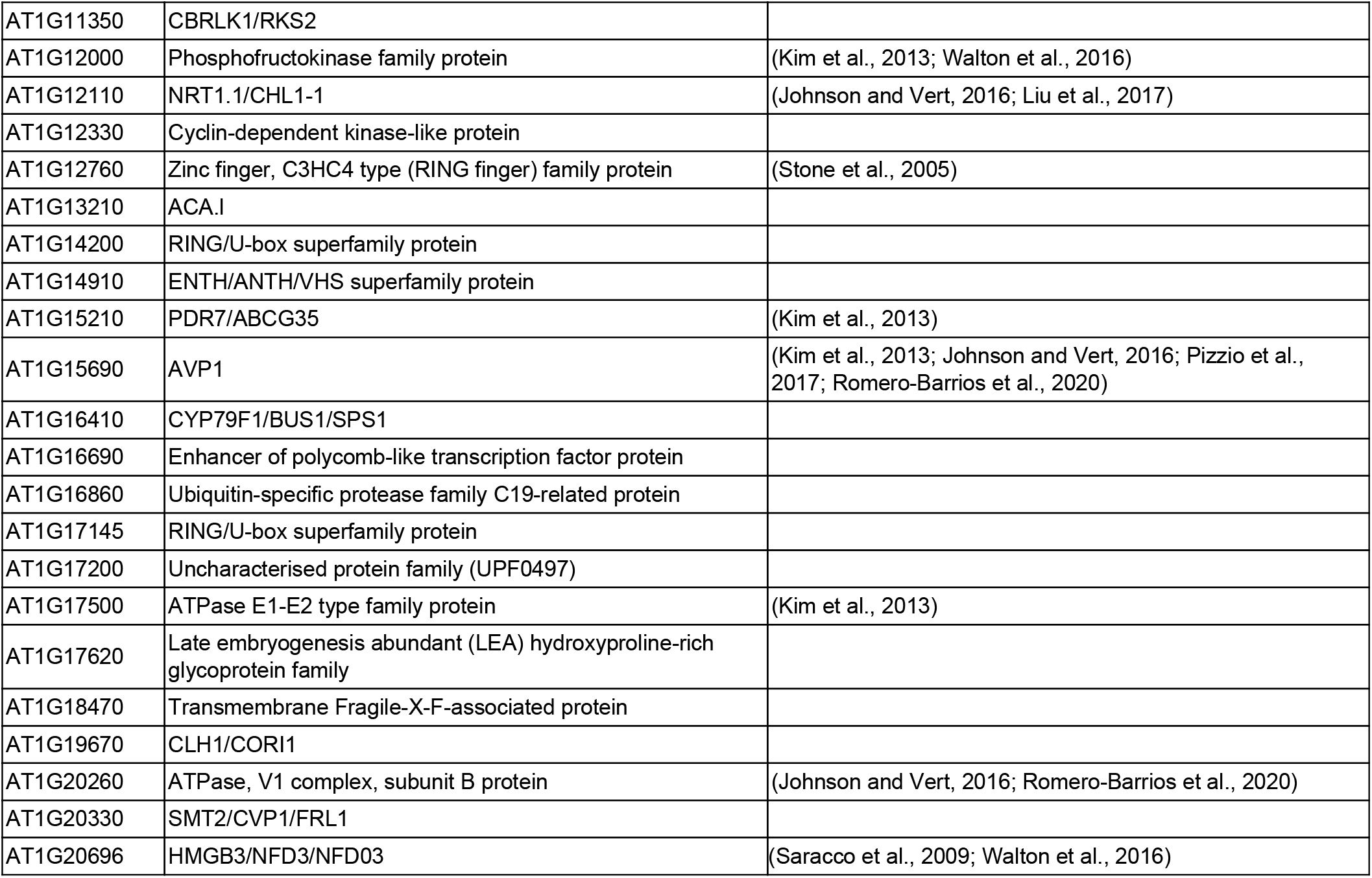
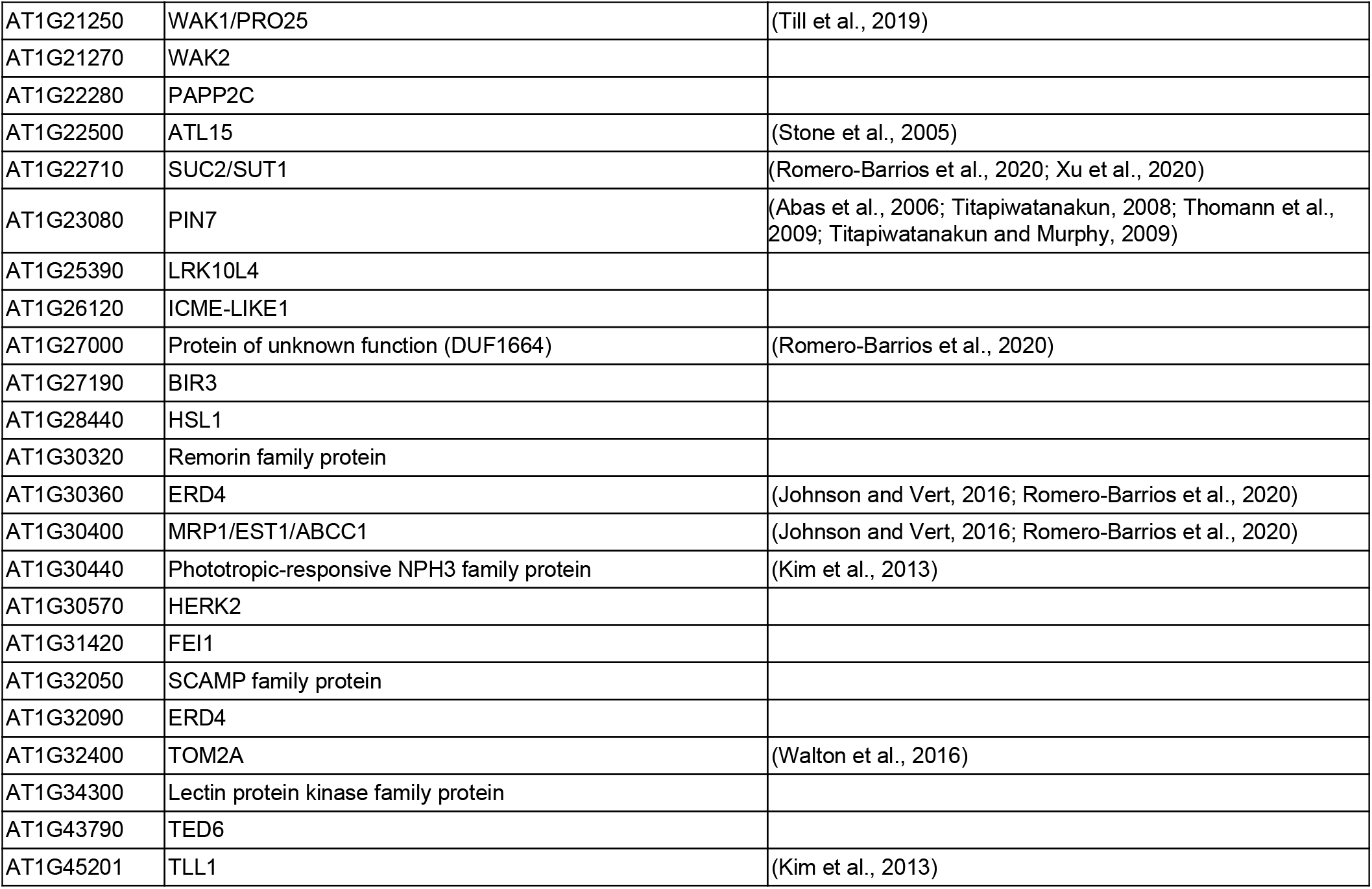
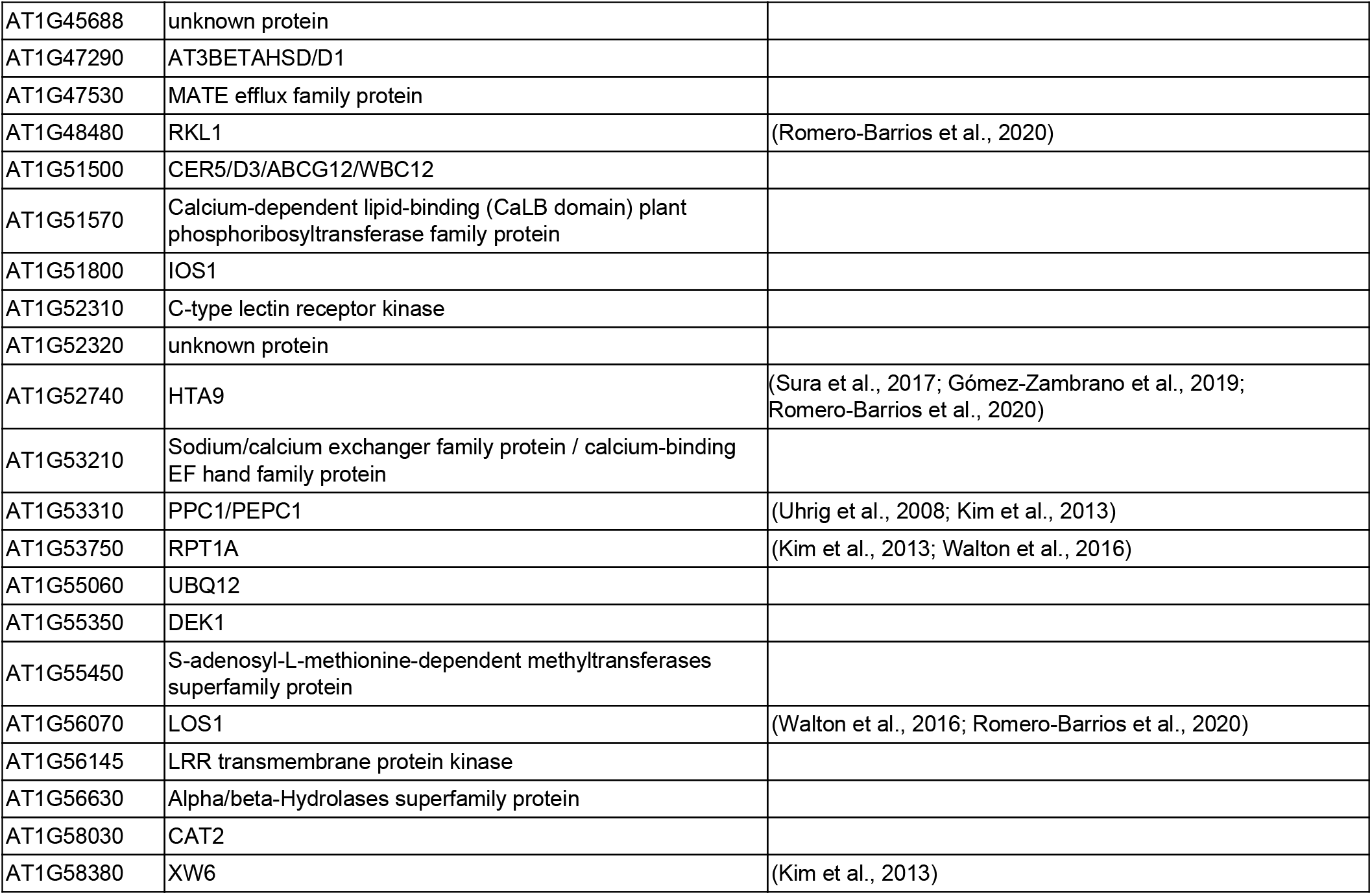
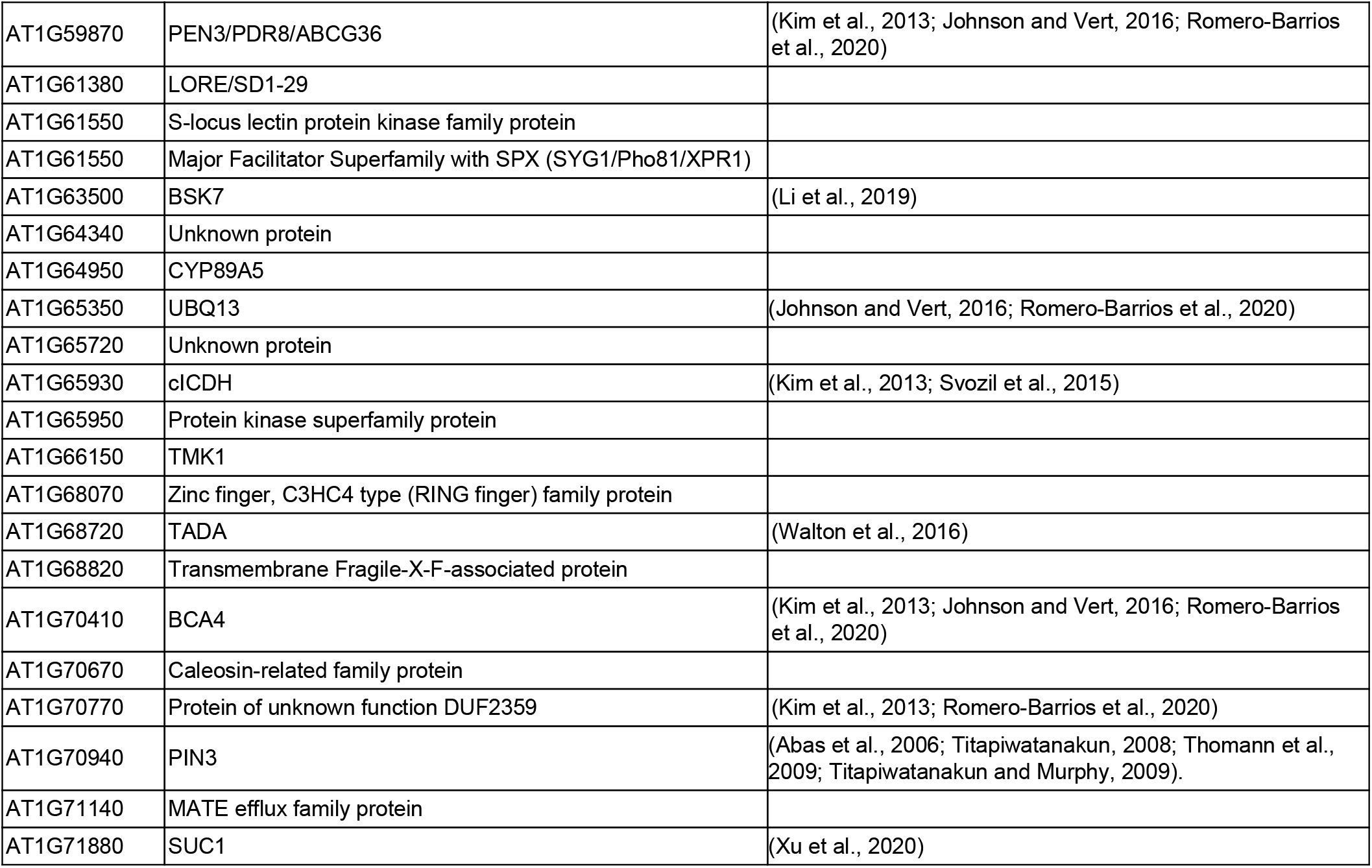
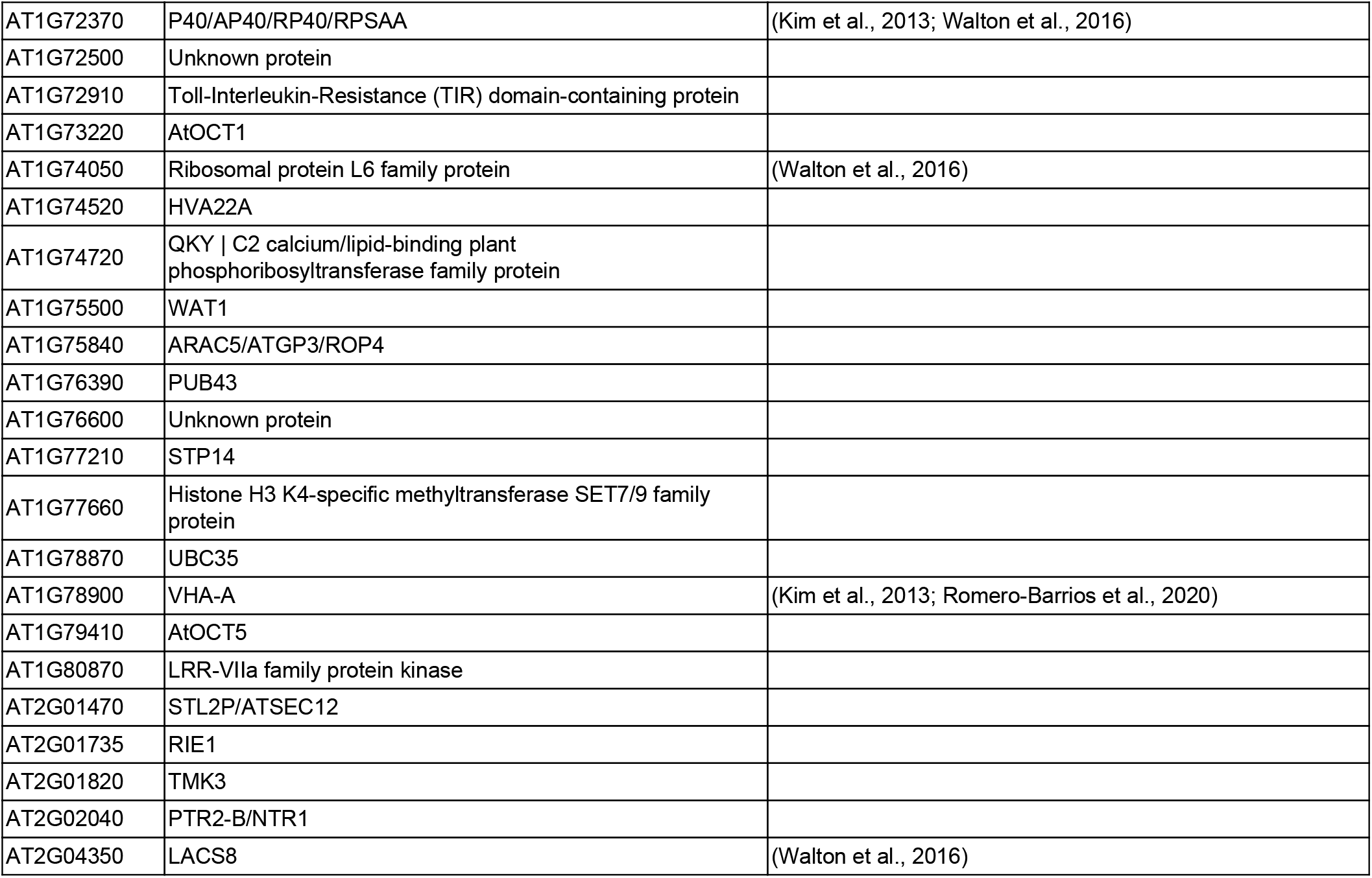
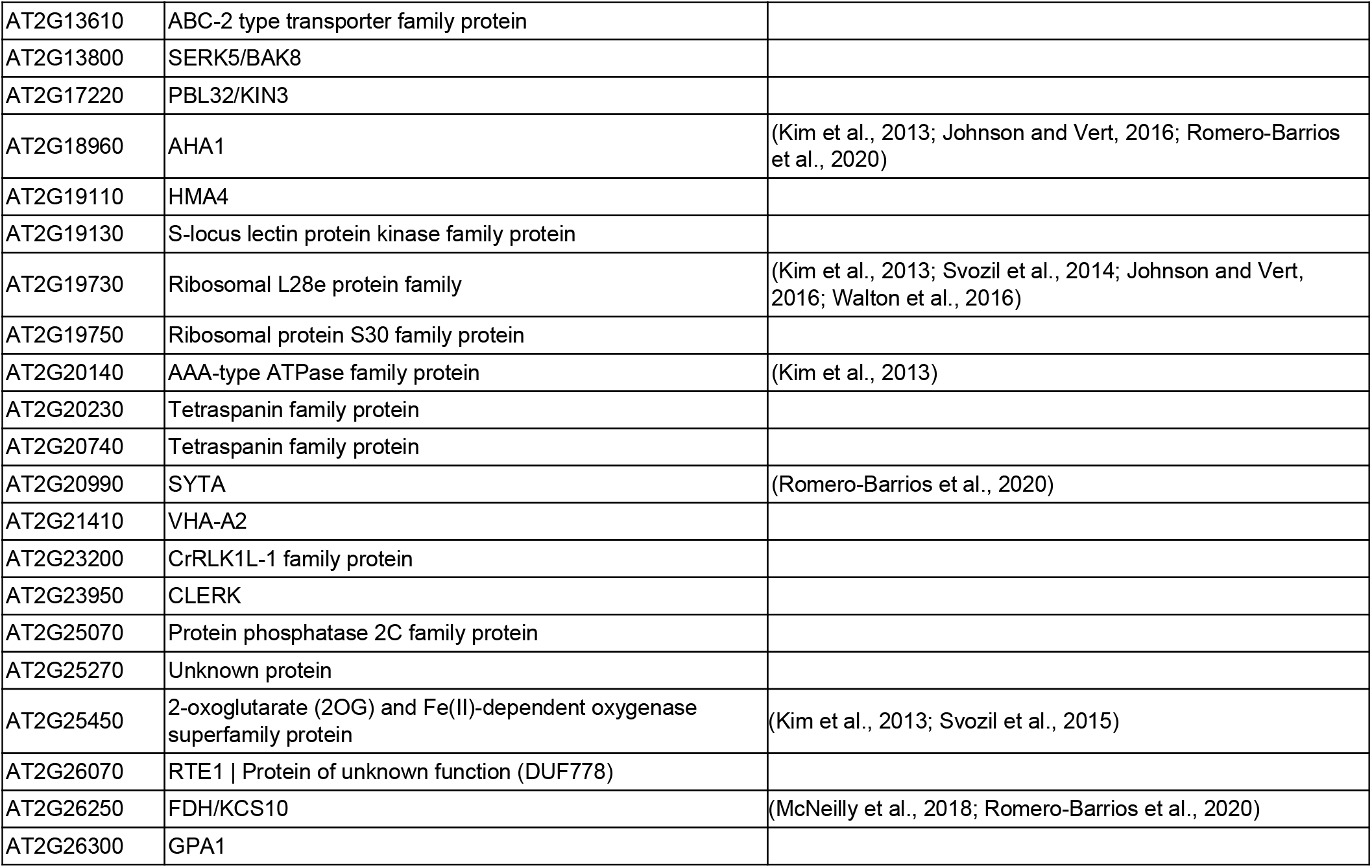
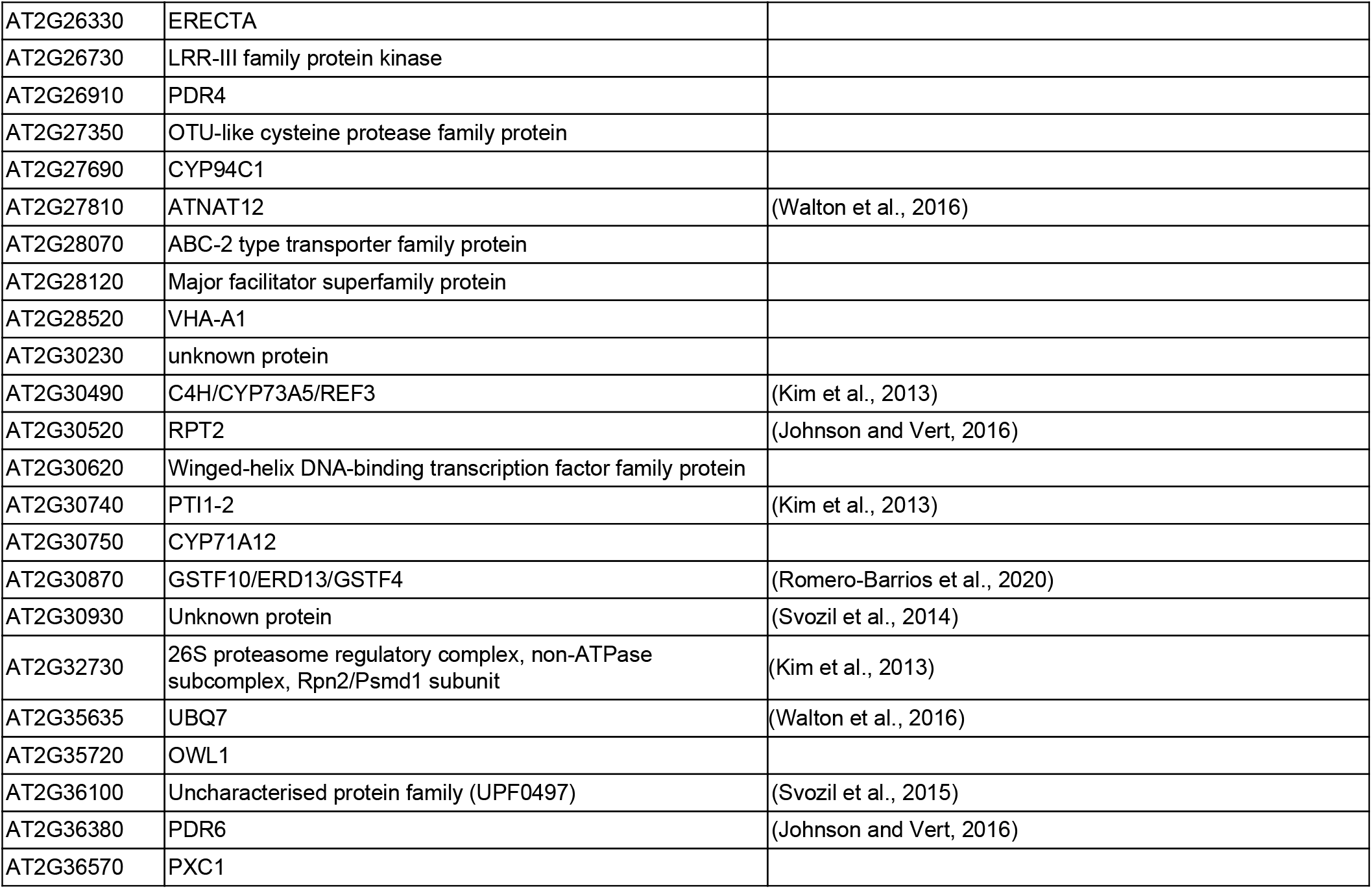
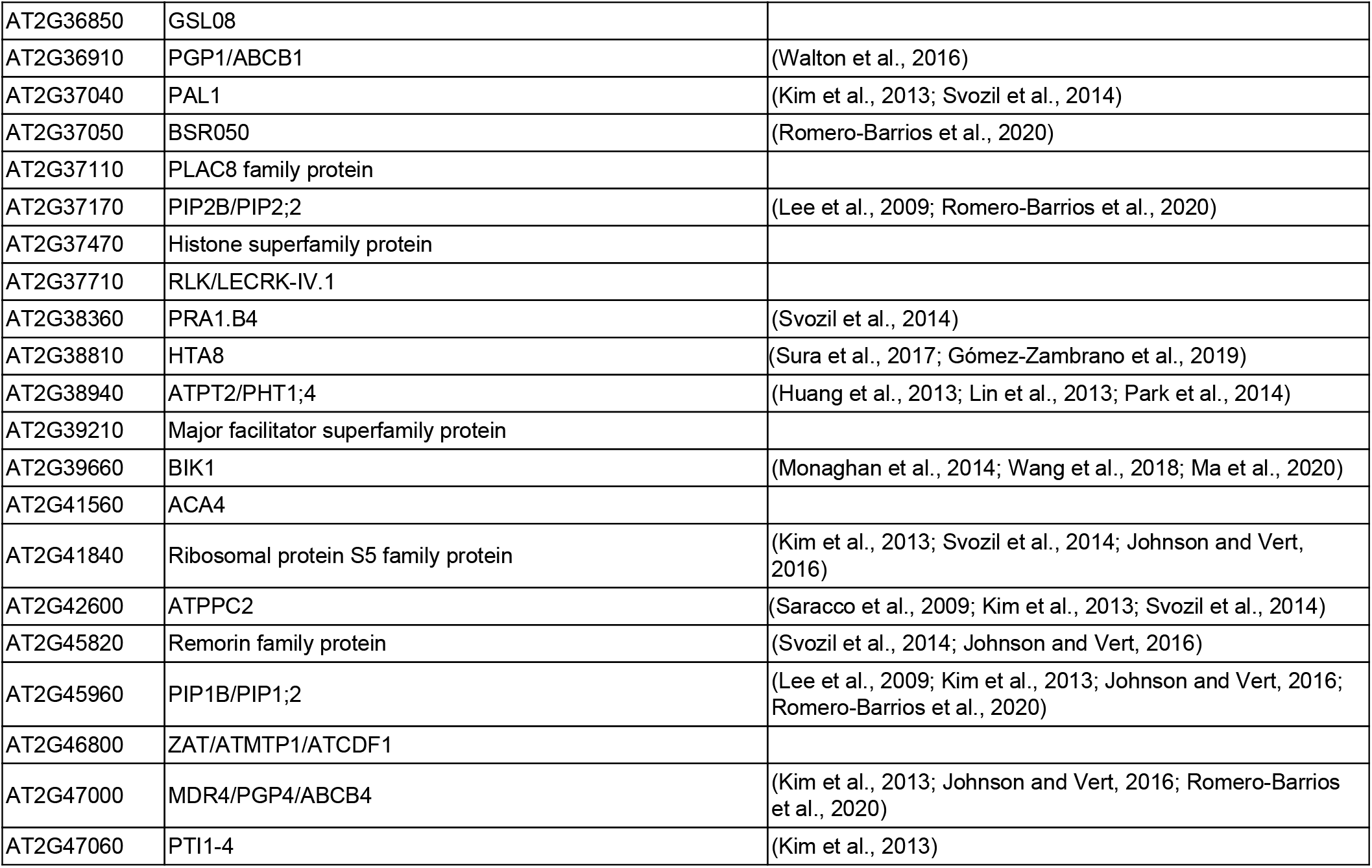
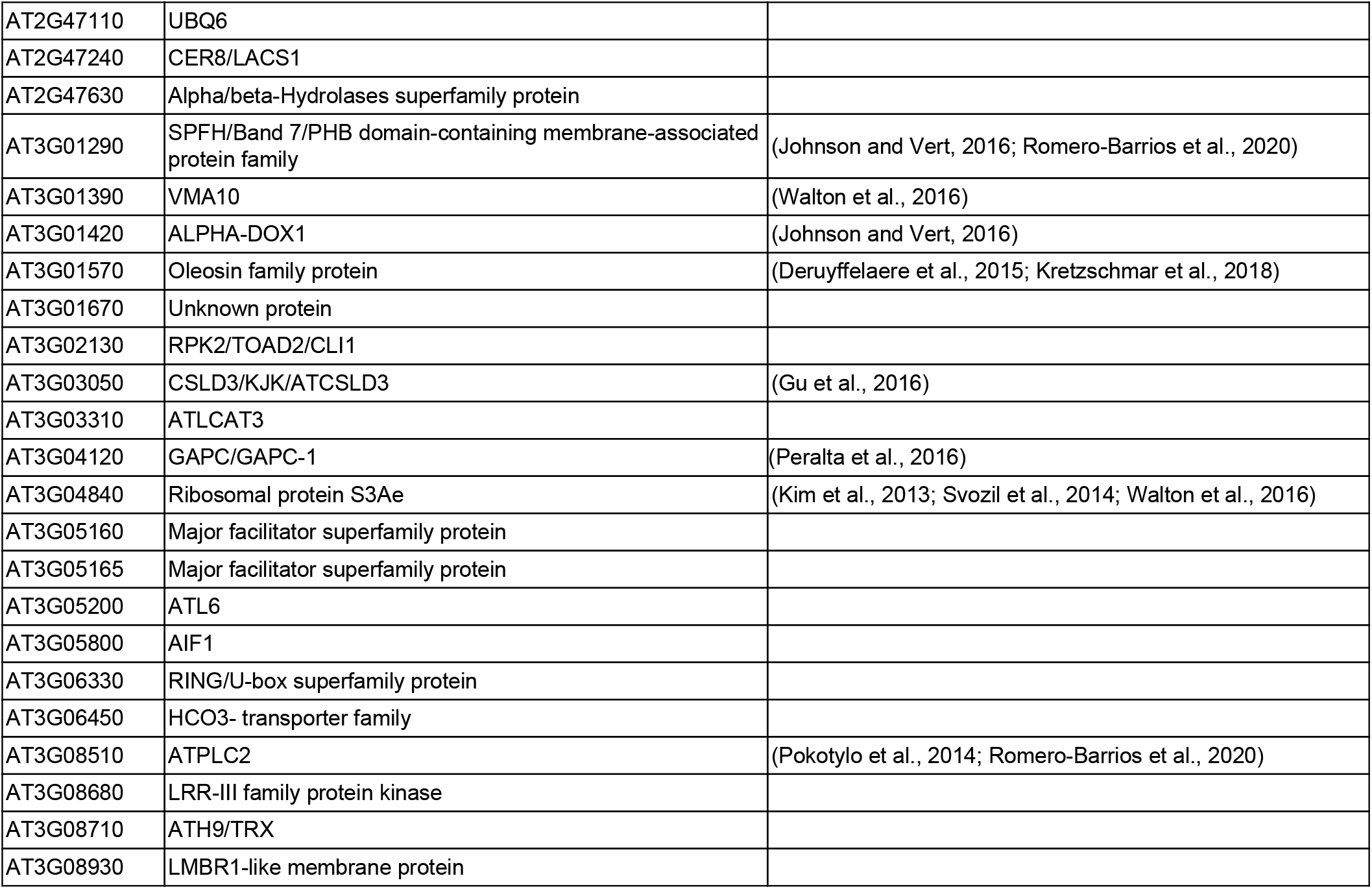
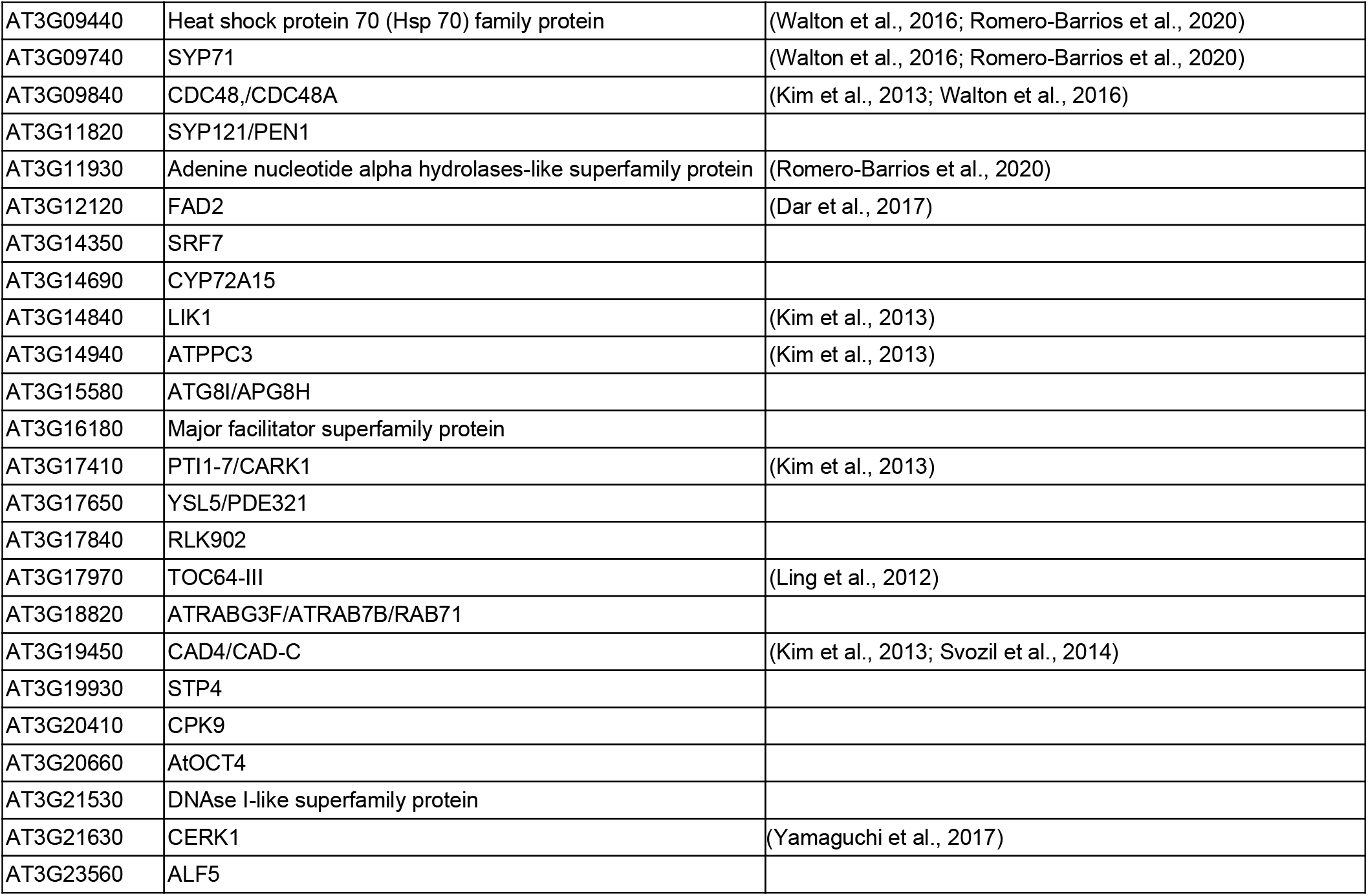
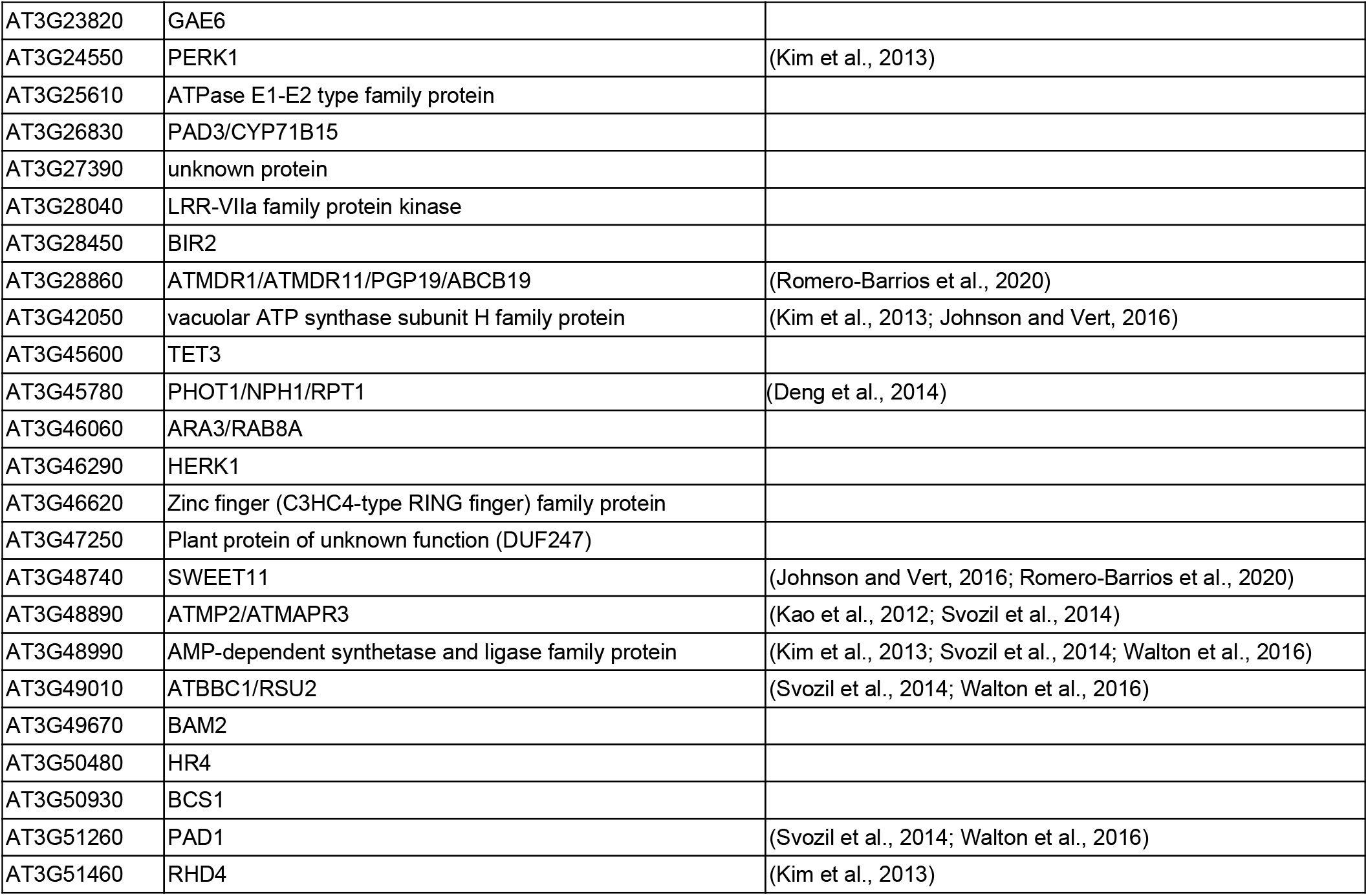
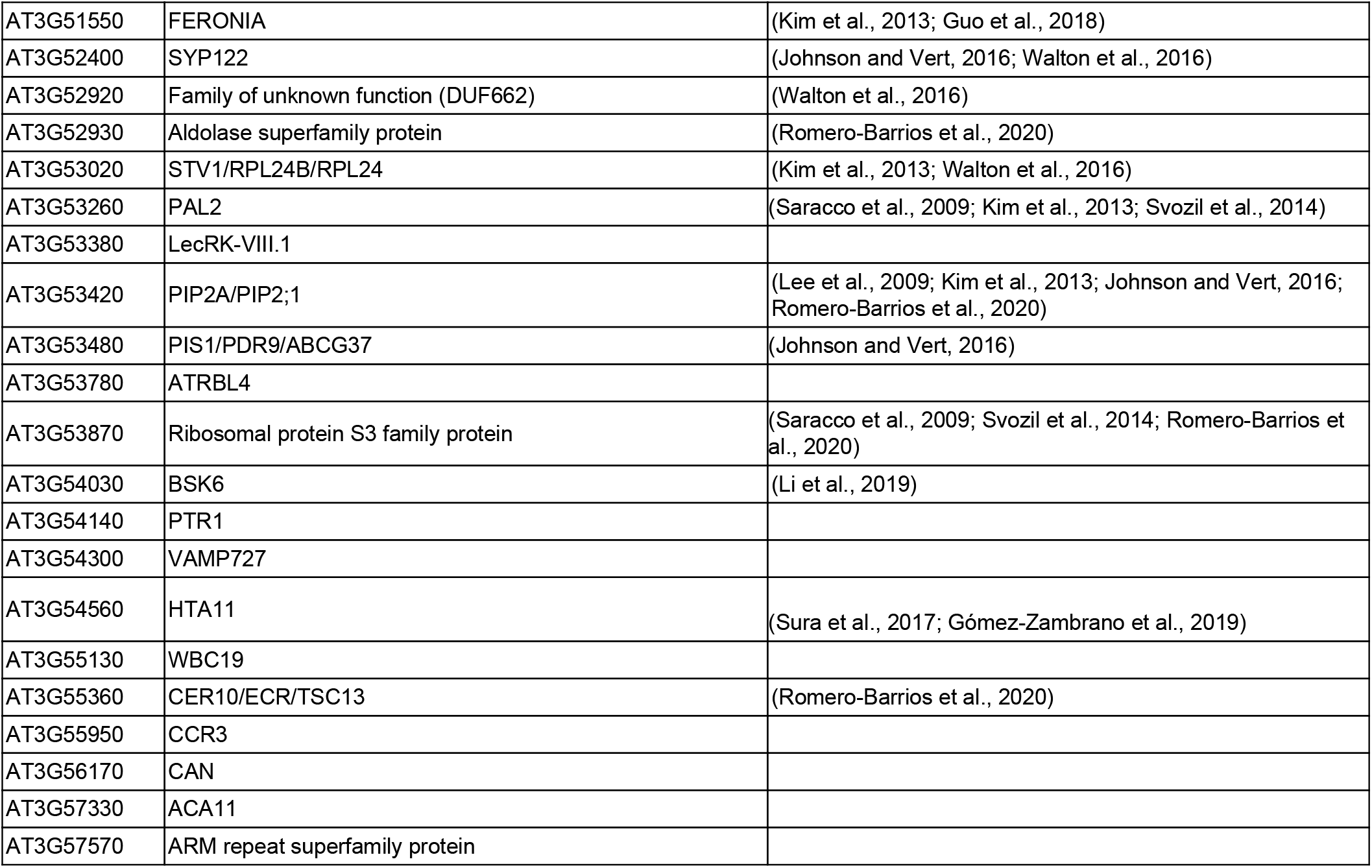
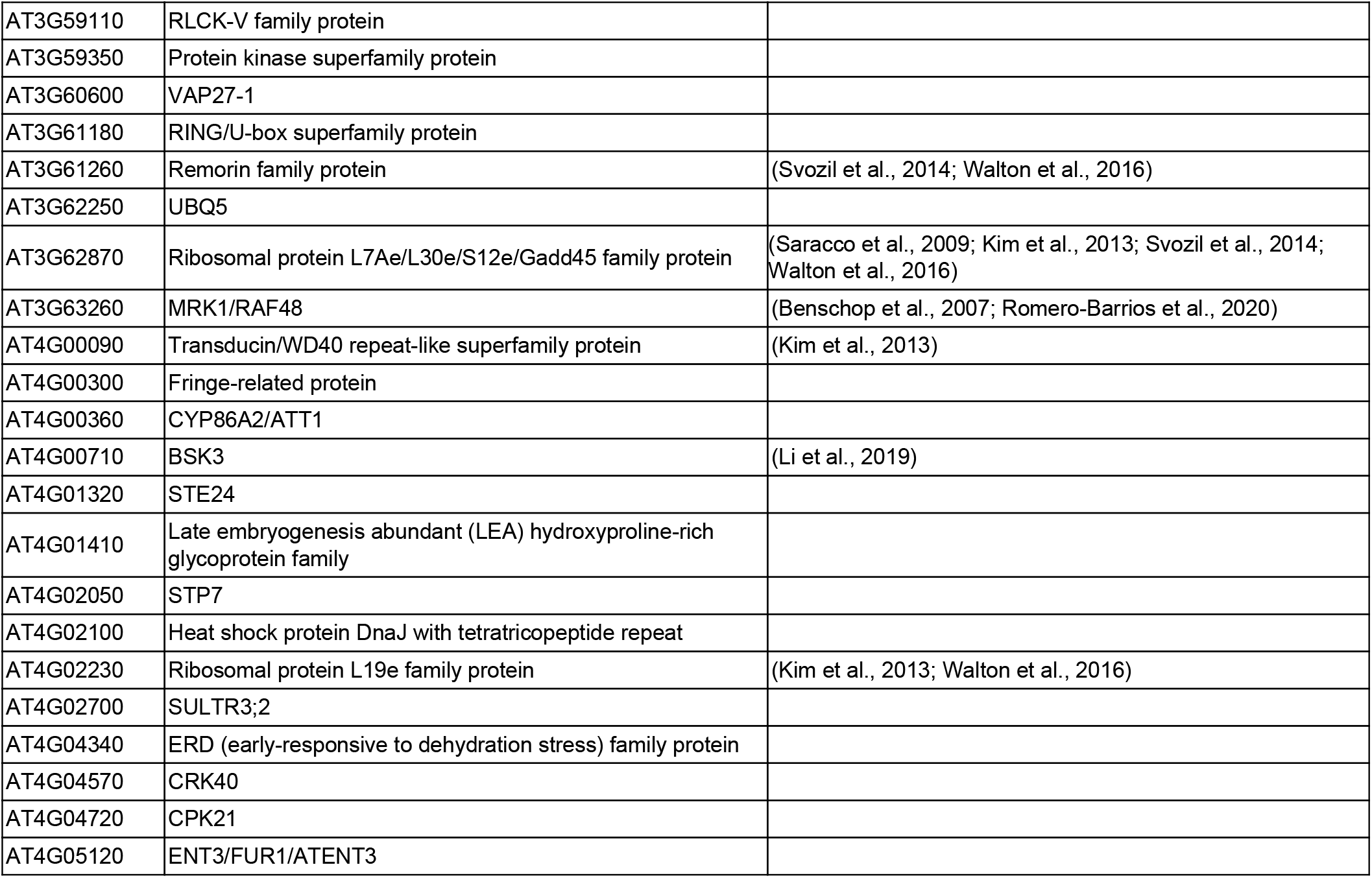
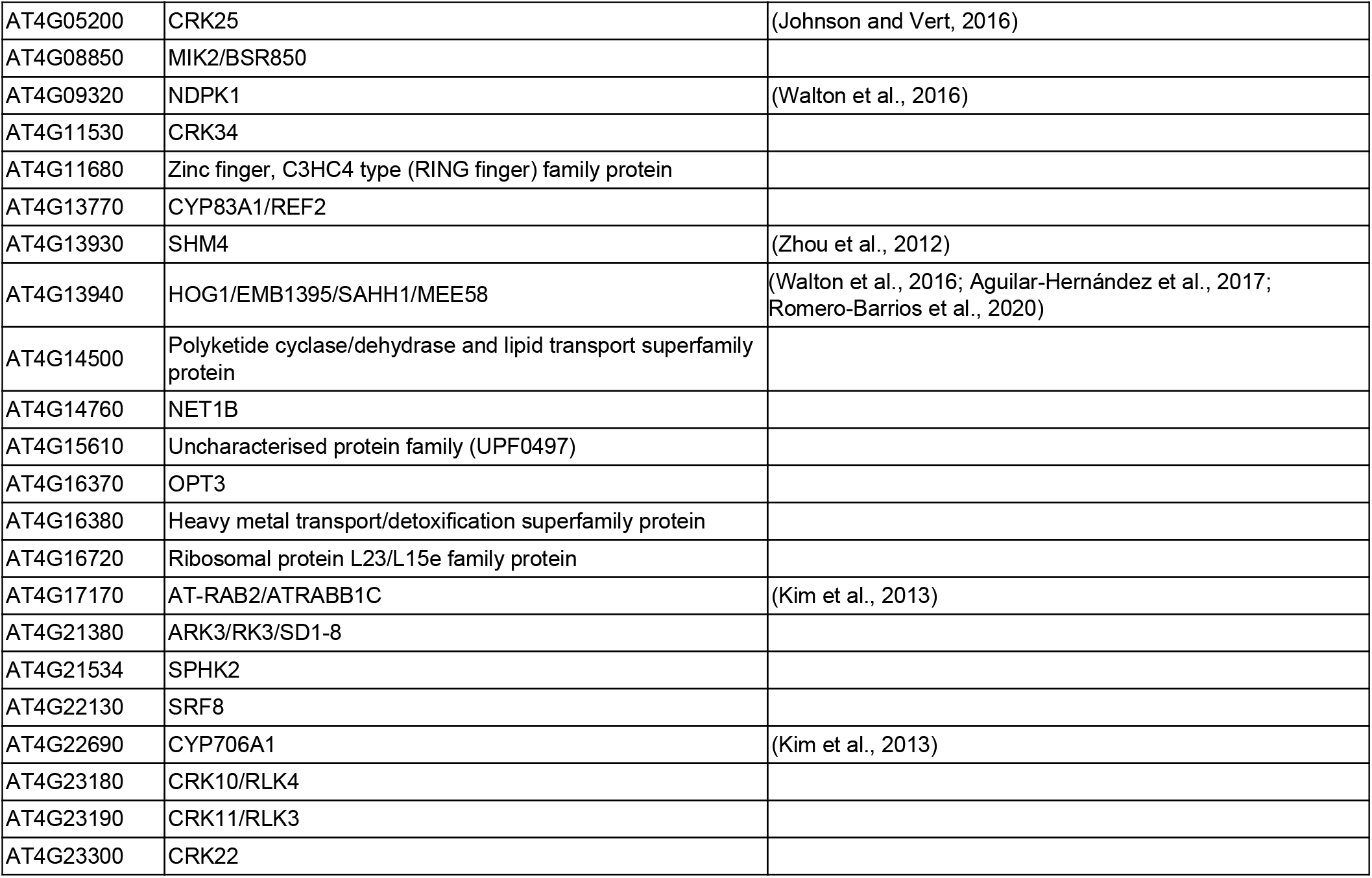
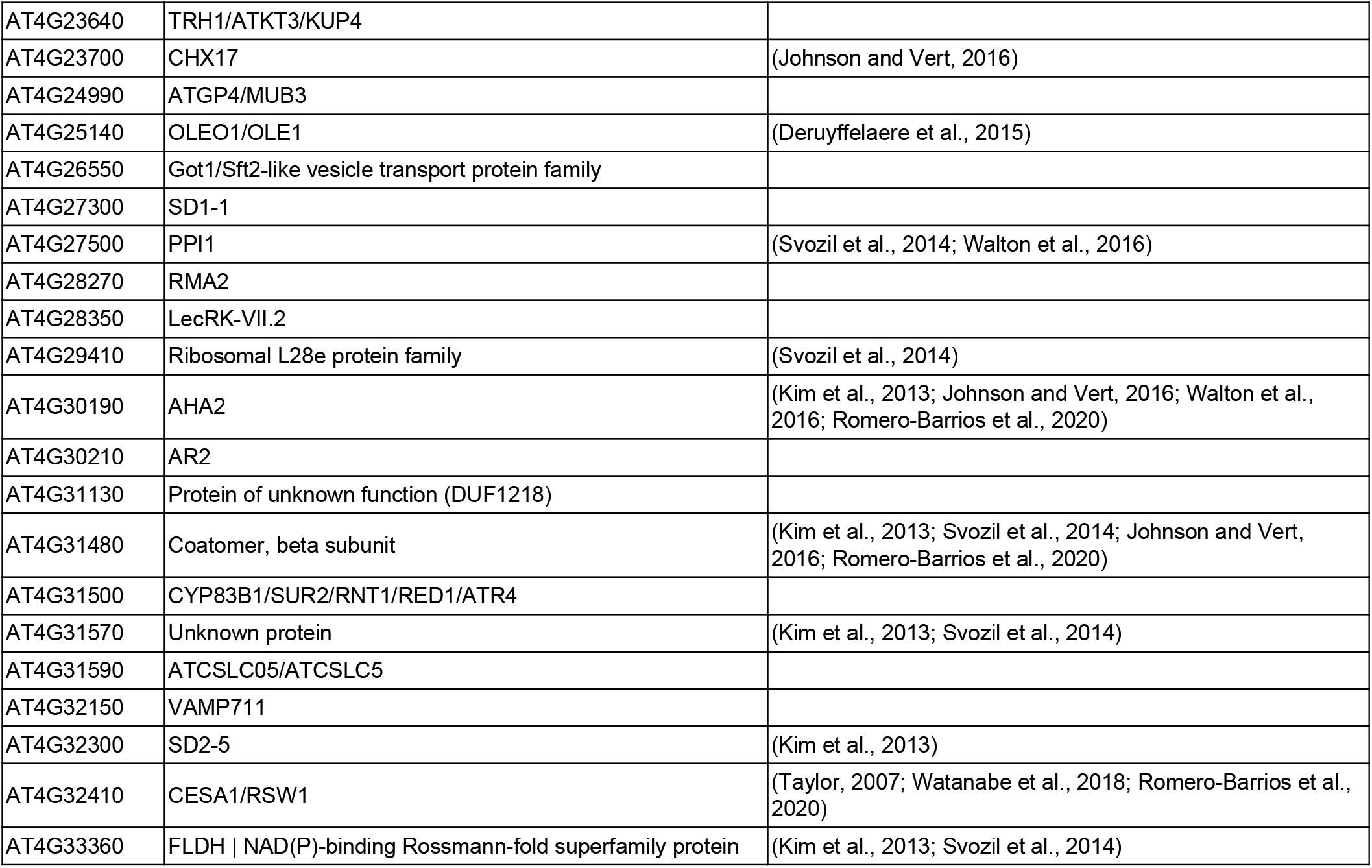
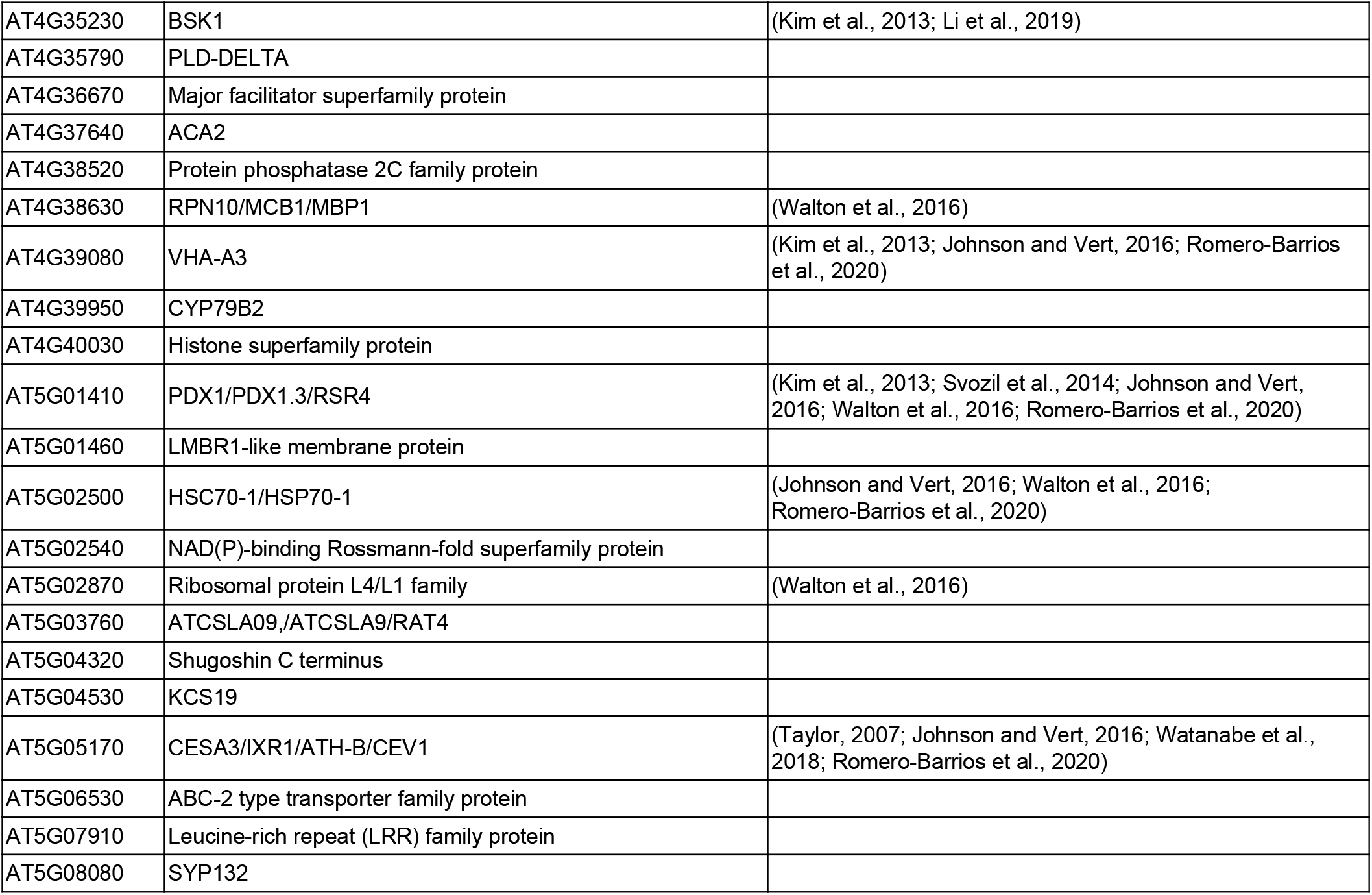
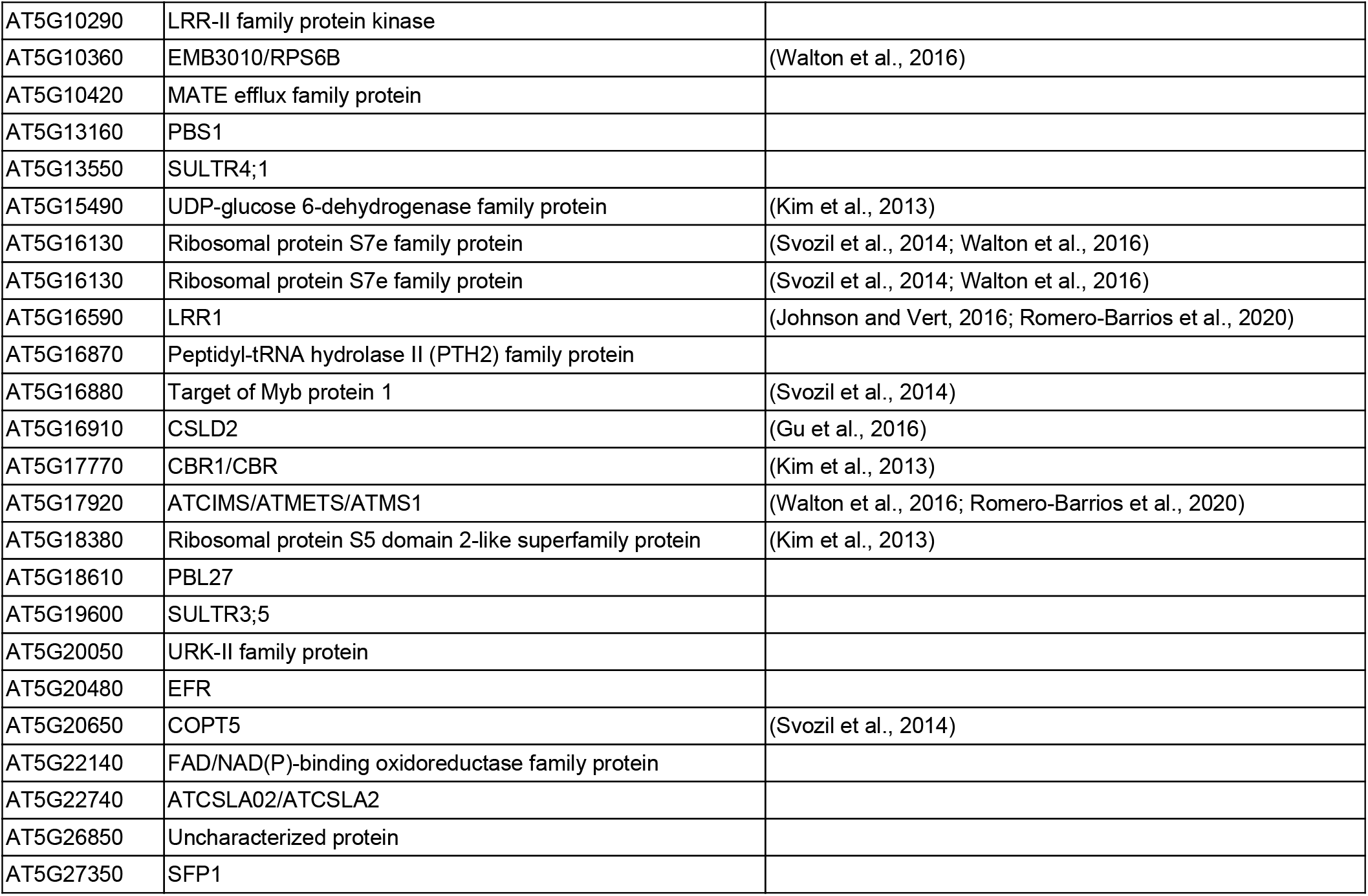
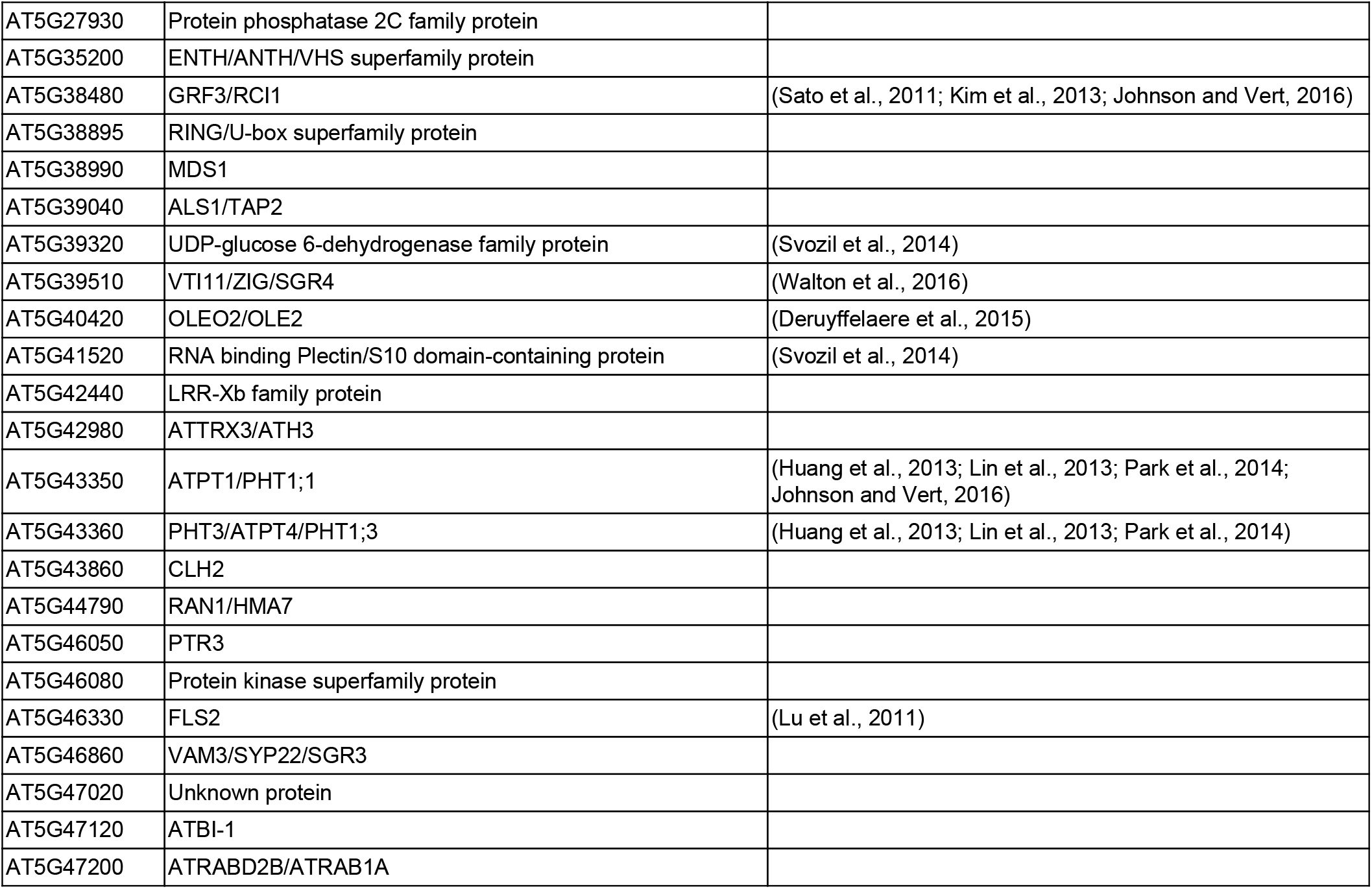
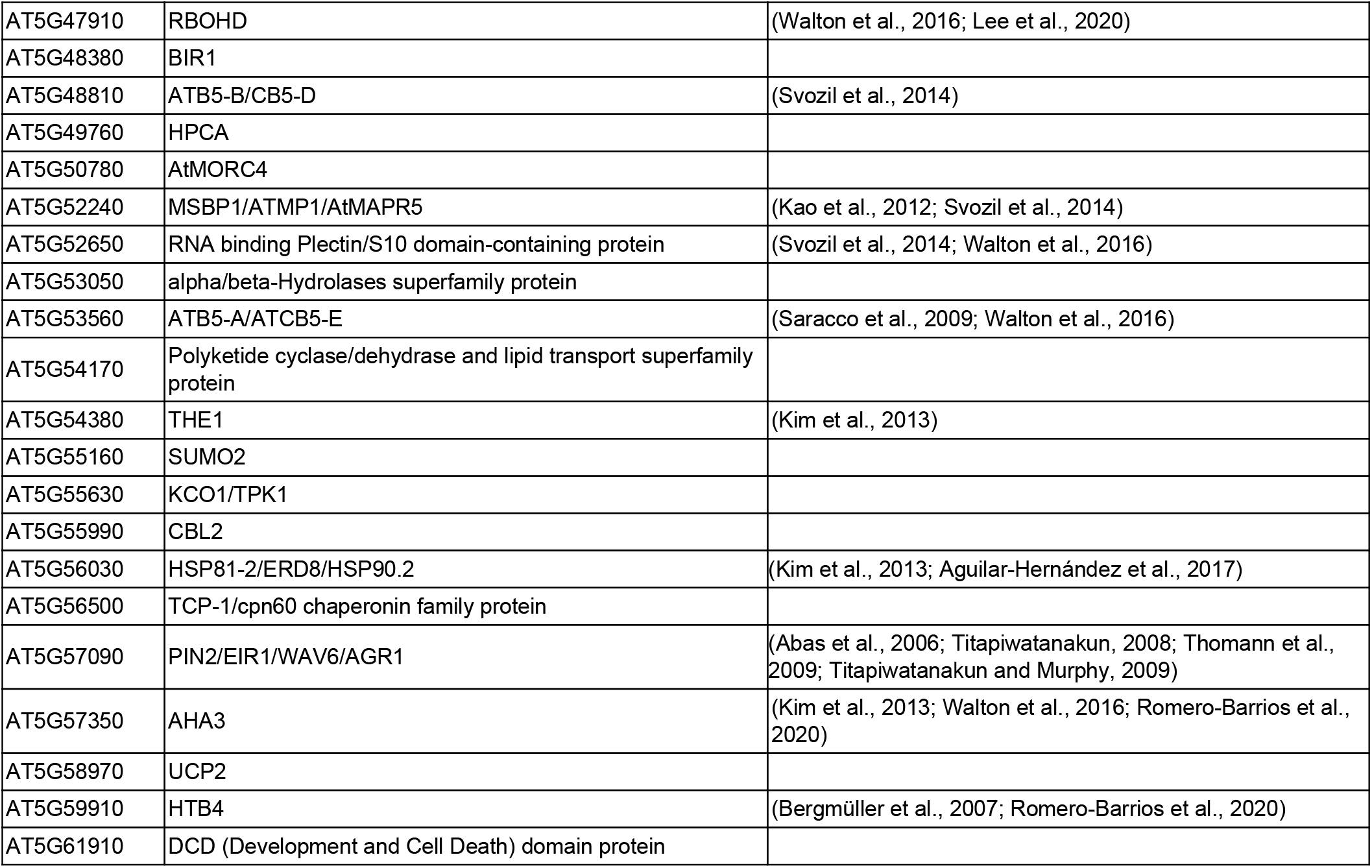
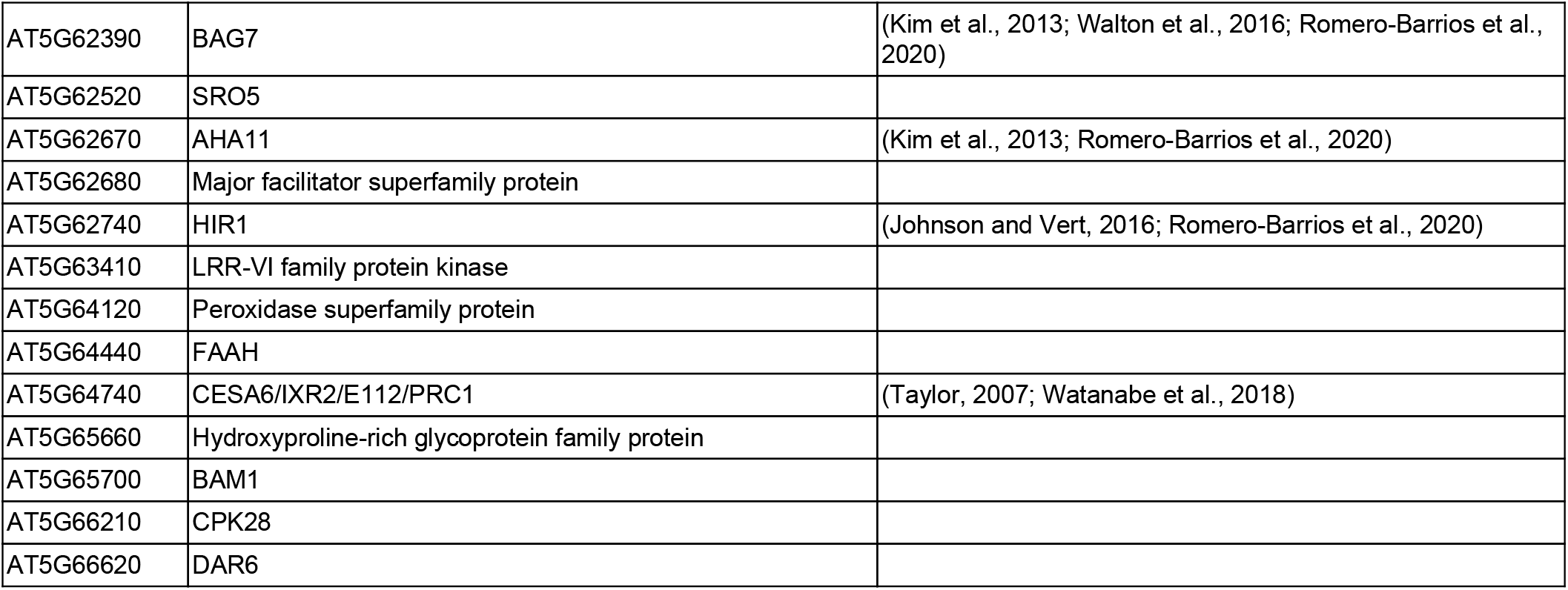
Comparative analysis reveals 265 unique ubiquitin targets identified in this study. Ubiquitinated proteins identified from earlier large-scale studies (Maor et al., 2007; Manzano et al., 2008; Igawa et al., 2009; Kim et al., 2013; Svozil et al., 2014; Johnson and Vert, 2016; Walton et al., 2016; Romero-Barrios et al., 2020) were compared to those identified in this study (Table S1) using Microsoft Excel. Manual inspection of the literature revealed additional ubiquitinated proteins, as well as proteins predicted to be ubiquitinated.

## References

Couto D, Niebergall R, Liang X, Bücherl CA, Sklenar J, Macho AP, Ntoukakis V, Derbyshire P, Altenbach D, Maclean D, et al (2016) The Arabidopsis Protein Phosphatase PP2C38 Negatively Regulates the Central Immune Kinase BIK1. PLoS Pathog 12: e1005811

Couto D, Zipfel C (2016) Regulation of pattern recognition receptor signalling in plants. Nat Rev Immunol 16: 537–552

Danielsen JMR, Sylvestersen KB, Bekker-Jensen S, Szklarczyk D, Poulsen JW, Horn H, Jensen LJ, Mailand N, Nielsen ML (2011) Mass spectrometric analysis of lysine ubiquitylation reveals promiscuity at site level. Mol Cell Proteomics 10: M110.003590

Jiang Y, Han B, Zhang H, Mariappan KG, Bigeard J, Colcombet J, Hirt H (2019) MAP4K4 associates with BIK1 to regulate plant innate immunity. EMBO Rep. 20:

Kelley LA, Mezulis S, Yates CM, Wass MN, Sternberg MJE (2015) The Phyre2 web portal for protein modeling, prediction and analysis. Nat Protoc 10: 845–858

Lal NK, Nagalakshmi U, Hurlburt NK, Flores R, Bak A, Sone P, Ma X, Song G, Walley J, Shan L, et al (2018) The Receptor-like Cytoplasmic Kinase BIK1 Localizes to the Nucleus and Regulates Defense Hormone Expression during Plant Innate Immunity. Cell Host Microbe 23: 485–497.e5

Liang X, Zhou J-M (2018) Receptor-Like Cytoplasmic Kinases: Central Players in Plant Receptor Kinase-Mediated Signaling. Annu Rev Plant Biol 69: 267–299

Ma X, Claus LAN, Leslie ME, Tao K, Wu Z, Liu J, Yu X, Li B, Zhou J, Savatin DV, et al (2020) Ligand-induced monoubiquitination of BIK1 regulates plant immunity. Nature 581: 199–203

Mithoe SC, Menke FL (2018) Regulation of pattern recognition receptor signalling by phosphorylation and ubiquitination. Curr Opin Plant Biol 45: 162–170

Monaghan J, Matschi S, Shorinola O, Rovenich H, Matei A, Segonzac C, Malinovsky FG, Rathjen JP, MacLean D, Romeis T, et al (2014) The calcium-dependent protein kinase CPK28 buffers plant immunity and regulates BIK1 turnover. Cell Host Microbe 16: 605–615

Paez Valencia J, Goodman K, Otegui MS (2016) Endocytosis and Endosomal Trafficking in Plants. Annu Rev Plant Biol 67: 309–335

Shiu S-H, Bleecker AB (2001) Plant Receptor-Like Kinase Gene Family: Diversity, Function, and Signaling. Science Signaling 2001: re22–re22

Shiu SH, Bleecker AB (2003) Expansion of the receptor-like kinase/Pelle gene family and receptor-like proteins in Arabidopsis. Plant Physiol 132: 530–543

Turek I, Tischer N, Lassig R, Trujillo M (2018) Multi-tiered pairing selectivity between E2 ubiquitin–conjugating enzymes and E3 ligases. J. Biol. Chem.

Udeshi ND, Mertins P, Svinkina T, Carr SA (2013) Large-scale identification of ubiquitination sites by mass spectrometry. Nat Protoc 8: 1950–1960

Vierstra RD (2012) The expanding universe of ubiquitin and ubiquitin-like modifiers. Plant Physiol 160: 2–14

Walton A, Stes E, Cybulski N, Van Bel M, Inigo S (2016) It’s time for some “site”-seeing: novel tools to monitor the ubiquitin landscape in Arabidopsis thaliana. The Plant

Wang J, Grubb LE, Wang J, Liang X, Li L, Gao C, Ma M, Feng F, Li M, Li L, et al (2018) A Regulatory Module Controlling Homeostasis of a Plant Immune Kinase. Mol Cell 69: 493–504.e6

Zhang M, Chiang Y-H, Toruño TY, Lee D, Ma M, Liang X, Lal NK, Lemos M, Lu Y-J, Ma S, et al (2018) The MAP4 Kinase SIK1 Ensures Robust Extracellular ROS Burst and Antibacterial Immunity in Plants. Cell Host Microbe 24: 379–391.e5

Zipfel C, Kunze G, Chinchilla D, Caniard A, Jones JDG, Boller T, Felix G (2006) Perception of the bacterial PAMP EF-Tu by the receptor EFR restricts Agrobacterium-mediated transformation. Cell 125: 749–760

## Supplemental References

Bailey TL, Boden M, Buske FA, Frith M, Grant CE, Clementi L, Ren J, Li WW, Noble WS (2009) MEME SUITE: tools for motif discovery and searching. Nucleic Acids Res 37: W202–8

Bender KW, Blackburn RK, Monaghan J, Derbyshire P, Menke FLH, Zipfel C, Goshe MB, Zielinski RE, Huber SC (2017) Autophosphorylation-based Calcium (Ca2+) Sensitivity Priming and Ca2+/Calmodulin Inhibition of Arabidopsis thaliana Ca2+-dependent Protein Kinase 28 (CPK28). J Biol Chem 292: 3988–4002

Crooks GE, Hon G, Chandonia J-M, Brenner SE (2004) WebLogo: a sequence logo generator. Genome Res 14: 1188–1190

Igawa T, Fujiwara M, Takahashi H, Sawasaki T, Endo Y, Seki M, Shinozaki K, Fukao Y, Yanagawa Y (2009) Isolation and identification of ubiquitin-related proteins from Arabidopsis seedlings. J Exp Bot 60: 3067–3073

Johnson A, Vert G (2016) Unraveling K63 Polyubiquitination Networks by Sensor-Based Proteomics. Plant Physiol 171: 1808–1820

Kim D-Y, Scalf M, Smith LM, Vierstra RD (2013) Advanced proteomic analyses yield a deep catalog of ubiquitylation targets in Arabidopsis. Plant Cell 25: 1523–1540

Manzano C, Abraham Z, López-Torrejón G, Del Pozo JC (2008) Identification of ubiquitinated proteins in Arabidopsis. Plant Mol Biol 68: 145–158

Maor R, Jones A, Nühse TS, Studholme DJ, Peck SC, Shirasu K (2007) Multidimensional protein identification technology (MudPIT) analysis of ubiquitinated proteins in plants. Mol Cell Proteomics 6: 601–610

Raudvere U, Kolberg L, Kuzmin I, Arak T, Adler P, Peterson H, Vilo J (2019) g:Profiler: a web server for functional enrichment analysis and conversions of gene lists (2019 update). Nucleic Acids Res 47: W191–W198

Romero-Barrios N, Monachello D, Dolde U, Wong A, San Clemente H, Cayrel A, Johnson A, Lurin C, Vert G (2020) Advanced Cataloging of Lysine-63 Polyubiquitin Networks by Genomic, Interactome, and Sensor-Based Proteomic Analyses. Plant Cell 32: 123–138

Svozil J, Hirsch-Hoffmann M, Dudler R, Gruissem W, Baerenfaller K (2014) Protein abundance changes and ubiquitylation targets identified after inhibition of the proteasome with syringolin A. Mol Cell Proteomics 13: 1523–1536

## Supplemental Table S4 References

Abas L, Benjamins R, Malenica N, Paciorek T, Wiśniewska J, Moulinier-Anzola JC, Sieberer T, Friml J, Luschnig C (2006) Intracellular trafficking and proteolysis of the Arabidopsis auxin-efflux facilitator PIN2 are involved in root gravitropism. Nat Cell Biol 8: 249–256

Aguilar-Hernández V, Kim D-Y, Stankey RJ, Scalf M, Smith LM, Vierstra RD (2017) Mass Spectrometric Analyses Reveal a Central Role for Ubiquitylation in Remodeling the Arabidopsis Proteome during Photomorphogenesis. Mol Plant 10: 846–865

Benschop JJ, Mohammed S, O’Flaherty M, Heck AJR, Slijper M, Menke FLH (2007) Quantitative phosphoproteomics of early elicitor signaling in Arabidopsis. Mol Cell Proteomics 6: 1198–1214

Bergmüller E, Gehrig PM, Gruissem W (2007) Characterization of post-translational modifications of histone H2B-variants isolated from Arabidopsis thaliana. J Proteome Res 6: 3655–3668

Dar AA, Choudhury AR, Kancharla PK, Arumugam N (2017) The FAD2 Gene in Plants: Occurrence, Regulation, and Role. Front Plant Sci 8: 1789

Deng Z, Oses-Prieto JA, Kutschera U, Tseng T-S, Hao L, Burlingame AL, Wang Z-Y, Briggs WR (2014) Blue light-induced proteomic changes in etiolated Arabidopsis seedlings. J Proteome Res 13: 2524–2533

Deruyffelaere C, Bouchez I, Morin H, Guillot A, Miquel M, Froissard M, Chardot T, D’Andrea S (2015) Ubiquitin-Mediated Proteasomal Degradation of Oleosins is Involved in Oil Body Mobilization During Post-Germinative Seedling Growth in Arabidopsis. Plant Cell Physiol 56: 1374–1387

Gómez-Zambrano Á, Merini W, Calonje M (2019) The repressive role of Arabidopsis H2A.Z in transcriptional regulation depends on AtBMI1 activity. Nature Communications. doi: 10.1038/s41467-019-10773-1

Gu F, Bringmann M, Combs J, Yang J, Bergmann D, Nielsen E (2016) The Arabidopsis CSLD5 functions in cell plate formation in a cell cycle dependent manner. The Plant Cell tpc.00203.2016

Guo H, Nolan TM, Song G, Liu S, Xie Z, Chen J, Schnable PS, Walley JW, Yin Y (2018) FERONIA Receptor Kinase Contributes to Plant Immunity by Suppressing Jasmonic Acid Signaling in Arabidopsis thaliana. Curr Biol 28: 3316–3324.e6

Huang T-K, Han C-L, Lin S-I, Chen Y-J, Tsai Y-C, Chen Y-R, Chen J-W, Lin W-Y, Chen P-M, Liu T-Y, et al (2013) Identification of downstream components of ubiquitin-conjugating enzyme PHOSPHATE2 by quantitative membrane proteomics in Arabidopsis roots. Plant Cell 25: 4044–4060

Kao A-L, Lin Y-H, Chen RP-Y, Huang Y-Y, Chen C-C, Yang C-C (2012) E3-independent ubiquitination of AtMAPR/MSBP1. Phytochemistry 78: 7–19

Kretzschmar FK, Mengel LA, Müller AO, Schmitt K, Blersch KF, Valerius O, Braus GH, Ischebeck T (2018) PUX10 Is a Lipid Droplet-Localized Scaffold Protein That Interacts with CELL DIVISION CYCLE48 and Is Involved in the Degradation of Lipid Droplet Proteins. Plant Cell 30: 2137–2160

Lee D, Lal NK, Lin Z-JD, Ma S, Liu J, Castro B, Toruño T, Dinesh-Kumar SP, Coaker G (2020) Regulation of reactive oxygen species during plant immunity through phosphorylation and ubiquitination of RBOHD. Nat Commun 11: 1838

Lee HK, Cho SK, Son O, Xu Z, Hwang I, Kim WT (2009) Drought Stress-Induced Rma1H1, a RING Membrane-Anchor E3 Ubiquitin Ligase Homolog, Regulates Aquaporin Levels via Ubiquitination in Transgenic Arabidopsis Plants. The Plant Cell 21: 622–641

Ling Q, Huang W, Baldwin A, Jarvis P (2012) Chloroplast biogenesis is regulated by direct action of the ubiquitin-proteasome system. Science 338: 655–659

Lin W-Y, Huang T-K, Chiou T-J (2013) NITROGEN LIMITATION ADAPTATION, a Target of MicroRNA827, Mediates Degradation of Plasma Membrane–Localized Phosphate Transporters to Maintain Phosphate Homeostasis in Arabidopsis. The Plant Cell 25: 4061–4074

Liu W, Sun Q, Wang K, Du Q, Li W-X (2017) Nitrogen Limitation Adaptation (NLA) is involved in source-to-sink remobilization of nitrate by mediating the degradation of NRT1.7 in Arabidopsis. New Phytologist 214: 734–744

Li Z, Shen J, Liang J (2019) Genome-Wide Identification, Expression Profile, and Alternative Splicing Analysis of the Brassinosteroid-Signaling Kinase (BSK) Family Genes in Arabidopsis. Int J Mol Sci. doi: 10.3390/ijms20051138

Lu D, Lin W, Gao X, Wu S, Cheng C, Avila J, Heese A, Devarenne TP, He P, Shan L (2011) Direct ubiquitination of pattern recognition receptor FLS2 attenuates plant innate immunity. Science 332: 1439–1442

McNeilly D, Schofield A, Stone SL (2018) Degradation of the stress-responsive enzyme formate dehydrogenase by the RING-type E3 ligase Keep on Going and the ubiquitin 26S proteasome system. Plant Mol Biol 96: 265–278

Park BS, Seo JS, Chua N-H (2014) NITROGEN LIMITATION ADAPTATION recruits PHOSPHATE2 to target the phosphate transporter PT2 for degradation during the regulation of Arabidopsis phosphate homeostasis. Plant Cell 26: 454–464

Peralta DA, Araya A, Busi MV, Gomez-Casati DF (2016) The E3 ubiquitin-ligase SEVEN IN ABSENTIA like 7 mono-ubiquitinates glyceraldehyde-3-phosphate dehydrogenase 1 isoform in vitro and is required for its nuclear localization in Arabidopsis thaliana. Int J Biochem Cell Biol 70: 48–56

Pizzio GA, Hirschi KD, Gaxiola RA (2017) Conjecture Regarding Posttranslational Modifications to the Arabidopsis Type I Proton-Pumping Pyrophosphatase (AVP1). Front Plant Sci 8: 1572

Pokotylo I, Kolesnikov Y, Kravets V, Zachowski A, Ruelland E (2014) Plant phosphoinositide-dependent phospholipases C: variations around a canonical theme. Biochimie 96: 144–157

Saracco SA, Hansson M, Scalf M, Walker JM, Smith LM, Vierstra RD (2009) Tandem affinity purification and mass spectrometric analysis of ubiquitylated proteins in Arabidopsis. Plant J 59: 344–358

Sato T, Maekawa S, Yasuda S, Domeki Y, Sueyoshi K, Fujiwara M, Fukao Y, Goto DB, Yamaguchi J (2011) Identification of 14-3-3 proteins as a target of ATL31 ubiquitin ligase, a regulator of the C/N response in Arabidopsis. The Plant Journal 68: 137–146

Stone SL, Hauksdóttir H, Troy A, Herschleb J, Kraft E, Callis J (2005) Functional analysis of the RING-type ubiquitin ligase family of Arabidopsis. Plant Physiol 137: 13–30

Sura W, Kabza M, Karlowski WM, Bieluszewski T, Kus-Slowinska M, Paweloszek L, Sadowski J, Ziolkowski PA (2017) Dual Role of the Histone Variant H2A.Z in Transcriptional Regulation of Stress-Response Genes. Plant Cell 29: 791–807

Svozil J, Gruissem W, Baerenfaller K (2015) Proteasome targeting of proteins in Arabidopsis leaf mesophyll, epidermal and vascular tissues. Front Plant Sci 6: 376

Taylor NG (2007) Identification of cellulose synthase AtCesA7 (IRX3) in vivo phosphorylation sites--a potential role in regulating protein degradation. Plant Mol Biol 64: 161–171

Thomann A, Lechner E, Hansen M, Dumbliauskas E, Parmentier Y, Kieber J, Scheres B, Genschik P (2009) Arabidopsis CULLIN3 Genes Regulate Primary Root Growth and Patterning by Ethylene-Dependent and -Independent Mechanisms. PLoS Genetics 5: e1000328

Till CJ, Vicente J, Zhang H, Oszvald M, Deery MJ, Pastor V, Lilley KS, Ray RV, Theodoulou FL, Holdsworth MJ (2019) The Arabidopsis thaliana N-recognin E3 ligase PROTEOLYSIS1 influences the immune response. Plant Direct 3: e00194

Titapiwatanakun B (2008) The regulation of P-glycoprotein and PIN by their cellular environment. Purdue University

Titapiwatanakun B, Murphy AS (2009) Post-transcriptional regulation of auxin transport proteins: cellular trafficking, protein phosphorylation, protein maturation, ubiquitination, and membrane composition. J Exp Bot 60: 1093–1107

Uhrig RG, She Y-M, Leach CA, Plaxton WC (2008) Regulatory monoubiquitination of phosphoenolpyruvate carboxylase in germinating castor oil seeds. J Biol Chem 283: 29650–29657

Watanabe Y, Schneider R, Barkwill S, Gonzales-Vigil E, Hill JL Jr, Samuels AL, Persson S, Mansfield SD (2018) Cellulose synthase complexes display distinct dynamic behaviors during xylem transdifferentiation. Proc Natl Acad Sci U S A 115: E6366–E6374

Xu Q, Yin S, Ma Y, Song M, Song Y, Mu S, Li Y, Liu X, Ren Y, Gao C, et al (2020) Carbon export from leaves is controlled via ubiquitination and phosphorylation of sucrose transporter SUC2. Proc Natl Acad Sci U S A 117: 6223–6230

Yamaguchi K, Mezaki H, Fujiwara M, Hara Y, Kawasaki T (2017) Arabidopsis ubiquitin ligase PUB12 interacts with and negatively regulates Chitin Elicitor Receptor Kinase 1 (CERK1). PLoS One 12: e0188886

Yun HS, Kwaaitaal M, Kato N, Yi C, Park S, Sato MH, Schulze-Lefert P, Kwon C (2013) Requirement of vesicle-associated membrane protein 721 and 722 for sustained growth during immune responses in Arabidopsis. Mol Cells 35: 481–488

Zhang N, Xu J, Liu X, Liang W, Xin M, Du J, Hu Z, Peng H, Guo W, Ni Z, et al (2019) Identification of HSP90C as a substrate of E3 ligase TaSAP5 through ubiquitylome profiling. Plant Sci 287: 110170

Zhou H, Zhao J, Yang Y, Chen C, Liu Y, Jin X, Chen L, Li X, Deng XW, Schumaker KS, et al (2012) UBIQUITIN-SPECIFIC PROTEASE16 Modulates Salt Tolerance in Arabidopsis by Regulating Na /H Antiport Activity and Serine Hydroxymethyltransferase Stability. The Plant Cell 24: 5106–5122

